# NTAD: A longitudinal neuroimaging dataset of Alzheimer’s disease pathology

**DOI:** 10.64898/2026.10.30.763020

**Authors:** Mats W.J. van Es, Marlou N. Perquin, Andrew J. Quinn, Jemma Pitt, Alexandra Krugliak, Tony Thayanandan, Ece Kocagoncu, Juliette Lanskey, Melek Karadag, Cara Alcock, Michelle Naessens, Stephen Lowe, Haddy Fye, Vanessa Raymont, Krish D. Singh, Richard N. Henson, Anna C. Nobre, Mark W. Woolrich, James B. Rowe, the NTAD study group

## Abstract

We introduce the New Therapeutics in Alzheimer’s Disease (NTAD) dataset, a multi-modal and longitudinal imaging dataset from 70 biomarker-positive people with early dementia or Mild Cognitive Impairment (MCI) due to Alzheimer’s Disease (AD), assessed over 2.5 years, and 35 healthy controls assessed at baseline. The data include simultaneous magneto-and electro-encephalography (M/EEG), 3-Tesla structural and functional magnetic resonance imaging (MRI), Amyloid-β positron emission tomography (PET), fluid proteomic biomarkers in cerebrospinal fluid and blood, and a battery of clinical and neuropsychological assessments. ^11^C UCB-J PET was undertaken for a subset of participants. For people with AD, assessments were repeated at approximately three weeks, 15 months, and 28 months after baseline. The data can be accessed through the Dementias Platform UK (DPUK), including pre-processed M/EEG and T1 MRI in pseudonymised BIDS format. With the pressing need to develop treatments for slowing, reversing, and preventing AD, these data provide an opportunity for identifying and validating novel biomarkers of Alzheimer’s disease neurophysiology, as well as understanding its treatable pathology.

## Background & Summary

Alzheimer’s Disease (AD) is the most common cause of dementia, with an estimated 50 million cases worldwide in 2020 and expected 150 million by 2050 (Patterson, 2020). While many disease-modifying treatments have been tested over the last decades, very few have demonstrated clinical success (Boxer & Sperling, 2023; Cummings et al., 2014; 2023; Mehta et al., 2018). One obstacle to successful trials is a lack of sensitive, robust markers of the multiple pathophysiological processes involved in AD. Such markers are essential to measure disease progression and treatment effects (e.g., Cacabelos, 2022; Kim et al., 2022). For biomarkers to be clinically informative and suitable for use in trials, they must be validated across three key characteristics: 1) reliability across repeated measurements, 2) sensitivity to the presence of AD, compared to healthy controls, and 3) sensitivity to longitudinal change over a timescale suitable for clinical trials. To achieve this, one needs evidence from baseline, test-retest, and longitudinal data.

Here we introduce the dataset from the New Therapeutics in Alzheimer’s Disease study (NTAD), a multi-modal and longitudinal dataset for investigating putative markers of AD. NTAD aims to validate neural markers recorded by magneto-/electro-encephalography (M/EEG). This data paper builds on a previously published protocol (Lanskey et al., 2022). A glossary of key terms and abbreviations can be found in Appendix A.

MEG may offer promising candidate markers for AD, as it captures neuronal population activity that is sensitive to changes in synaptic functioning. Synaptic dysfunction is an early feature of AD preceding larger characteristic structural changes (Busche & Hyman, 2020; DeKosky & Scheff, 1990; Selkoe, 2002; Terry et al., 1991). Differences in MEG resting state connectivity have been observed between healthy participants with amyloid-positive versus amyloid-negative status, even in the absence of metabolic or structural abnormalities (Nakamura et al., 2017). In clinical groups, MEG measures have been shown to capture disease related changes in resting state and task evoked activity in AD and other neurodegenerative conditions (Sitnikova et al., 2018; Puttaert et al., 2020; Cope et al., 2022; Kocagoncu et al., 2020; 2022; Sami et al., 2018; see Mandal et al., 2018 for a review). Multi-modal studies further relate M/EEG changes to amyloid and tau pathology, with evidence for nonlinear trajectories across disease stages, highlighting the need for longitudinal studies (Sohrabpour et al., 2025; also see Alfonsín et al., 2026). MEG is also sensitive to pharmacologically induced changes in neural activity, including GABAergic, glutamatergic, and serotonergic modulation, supporting its potential as an outcome measure in clinical trials (Adams et al., 2020; Hughes et al., 2015; 2025; Lanskey et al., 2025; Perry et al., 2022; Williams et al., 2025).

The NTAD study includes people with amyloid-positive AD and Mild Cognitive Impairment (AD/MCI), undertaking a battery of clinical assessments and neurophysiological tests. Amyloid positivity was established with cerebrospinal fluid (CSF) obtained by lumbar puncture (LP) and/or with positron emission tomography (PET) evidence of amyloid tracer retention in people meeting clinical diagnostic criteria of AD (Mckhann et al., 2011). Blood biomarkers of AD were performed retrospectively. Participants also completed about an hour of structural and functional magnetic resonance imaging (MRI), with sequences aligned by design to the Deep and Frequent Phenotyping (DFP) study of prodromal AD (Koychev et al., 2018). Estimates of synaptic density were measured in a subset of participants using ^11^C UCB-J PET, which quantifies synaptic vesicle glycoprotein 2A, SV2A, as an in vivo proxy of synaptic density (Nabulsi et al., 2016; O’Dell et al., 2021), and has shown sensitivity to AD and other dementias (Mecca et al., 2022; Carson et al., 2022; O’Dell et al., 2021; Holland et al., 2023; Malpetti et al., 2023). NTAD includes phlebotomy to extract DNA for genetic risk testing, as well as plasma, serum, and RNA aliquots. Analyses from a subset of the NTAD MEG data have been published previously (Jafarian et al., 2024; 2025; Lanskey et al., 2025a; 2025b; Trubshaw et al., 2026). A subset of processed blood samples contributed to a larger cohort study across multiple neurodegenerative diseases (Malpetti et al., 2025).

Various large scale open MEG datasets related to aging and neurodegeneration have been released in recent years, placing NTAD in a solid MEG translational landscape. The Cambridge Centre for Ageing and Neuroscience study (Cam CAN; phases 1 to 3; Shafto et al., 2014; Taylor et al., 2017) provides baseline data from approximately 700 healthy individuals across the adult lifespan, as well as 12-year follow up data on a subset of approximately 150 (phases 4 to 5; Demetriou et al., 2025). In addition, the “cognitively frail” sub study of Cam CAN includes approximately 80 older healthy participants, of whom half have poor cognitive performance but no subjective memory complaints (Kocagoncu et al., 2021). More lifespan data is available through the Welsh Advanced Neuroimaging Database (WAND; McNabb et al., 2025), including baseline MEG and other modalities in approximately 170 healthy volunteers across a wide age range, and the Open MEG Archive (OMEGA), including healthy as well as various disease populations (total N currently >600; Niso et al., 2016). Several databases focus on the prodromal stage of AD. For example, the “BioFIND” dataset contains resting state MEG data from 324 people, approximately half of whom had mild cognitive impairment (MCI; Hughes et al., 2019; Vaghari et al., 2022). The AI MIND cohort has studied ∼200 people with MCI using MEG, among the ∼1000 with EEG (www.ai-mind.eu). A smaller number of cohorts have longitudinal data, such as the Deep and Frequent Phenotyping study (DFP; Koychev et al., 2018), which includes repeat multi-modal, multi-site data using the same MEG and MRI acquisition protocols on approximately 100 participants with high (80%) and low risk (20%) of AD across multiple sites over the course of a year, aligned with the NTAD protocols. The PREVENT AD study, hosted through OMEGA, provides additional prodromal resting state data from approximately 100 participants of whom about 30 converted to MCI after a period of six years, including repeat MEG of individuals who do (N=20) and do not (N=50) convert (Villeneuve et al., 2025). NTAD extends these cohorts by measuring both resting state and task-based MEG longitudinally in individuals with AD or MCI over the course 0.5-2.5 years.

NTAD can be found on the DPUK Data Platform (Bauermeister et al., 2020). As the whole study dataset contains sensitive (clinical) patient information, the data can be accessed and processed on the data portal directly rather than by distribution of data copies. Personally identifiable data are not available via DPUK, for compliance with the General Data Protection Regulations, and access to data is conditional on not attempting to de-identify participants. M/EEG, PET, and MRI data are available in standardised Brain Imaging Data Structure (BIDS; Gorgolewski et al., 2016; Niso et al., 2018) format. Non-imaging data, including clinical and cognitive assessments, amyloid status, demographics, and metadata, are available in CSV format alongside the imaging data. For MEG, we share raw and preprocessed data. Further preprocessed data (including results from blood proteomics and standardized uptake value ratio (SUVR) for UCB-J PET) will be made available in future releases.

## Methods

### Participants

Participants were recruited between 2018 and 2022 from two UK sites and surrounding areas (Cambridge and Oxford, United Kingdom), including Cambridge University Hospitals NHS Trust memory clinics, Cambridge and Oxford research participation networks, the Cambridgeshire and Peterborough NHS Foundation Trust, regional memory clinics, and the NIHR Joint Dementia Research platforms. The study was approved by the East of England Cambridge Central Research Ethics Committee (REC reference 18/EE/0042), and participants gave written informed consent in accordance with the Declaration of Helsinki (1991). The study sought participants aged 50-85 (balanced numbers of male and female participants) with symptomatic MCI or early dementia due to Alzheimer’s disease (AD), and age-, sex-, and education-matched people with normal cognition. However, due to recruitment challenges, a strict balance in age and sex, as well as a matched population of people with normal cognition were not fully achieved (see below).

After pre-screening with medical records, initial postal contact and telephone discussion, 121 people with MCI/AD and 41 healthy controls were screened in person using a staged screening process to assess study eligibility (Figure 1), as outlined in detail in the NTAD protocol paper (Lanskey et al., 2022). In summary, patients were eligible if they had a Mini-Mental State Examination (MMSE) score above 18 and a Clinical Dementia Rating (CDR) of 0.5 or 1 in addition to a positive biomarker status, as assessed through amyloid positron emission tomography (PET) or cerebrospinal fluid (CSF) collection. Positive amyloid status was indicated by florbetaben scans with Standardized Uptake Value Ratio (SUVR) >1.19 (PET; Sabri et al., 2015; Rowe et al., 2017), or a total τ to amyloid β1-42 ratio >1 and amyloid β1-42 concentration <450 pg/mL or total phosphorylated τ >57 (CSF). Two additional participants were retained at the discretion of the lead physician: one did have not have SUVR classification so their PET images were visually assessed, and the other was a patient as the in-house clinic whose τ/amyloid β1-42 ratio was near threshold (.91). Healthy controls were eligible if their MMSE was above 24 and their CDR was 0. Their amyloid status was assessed through PET or CSF, but this was not used as an inclusion criterion. All participants were assessed by a physician during screening to ensure that cognitive impairments were not potentially caused by factors other than AD (e.g., stroke, trauma, medication effects), and that participants had sufficient health and no contraindications to complete the study procedures. All participants were required to bring a study partner, such as a spouse or family member, to the screening session. Study partners joined the medical interview and completed two questionnaires assessing the participant. Both participants and study partners signed informed consent.

**Figure 1.**
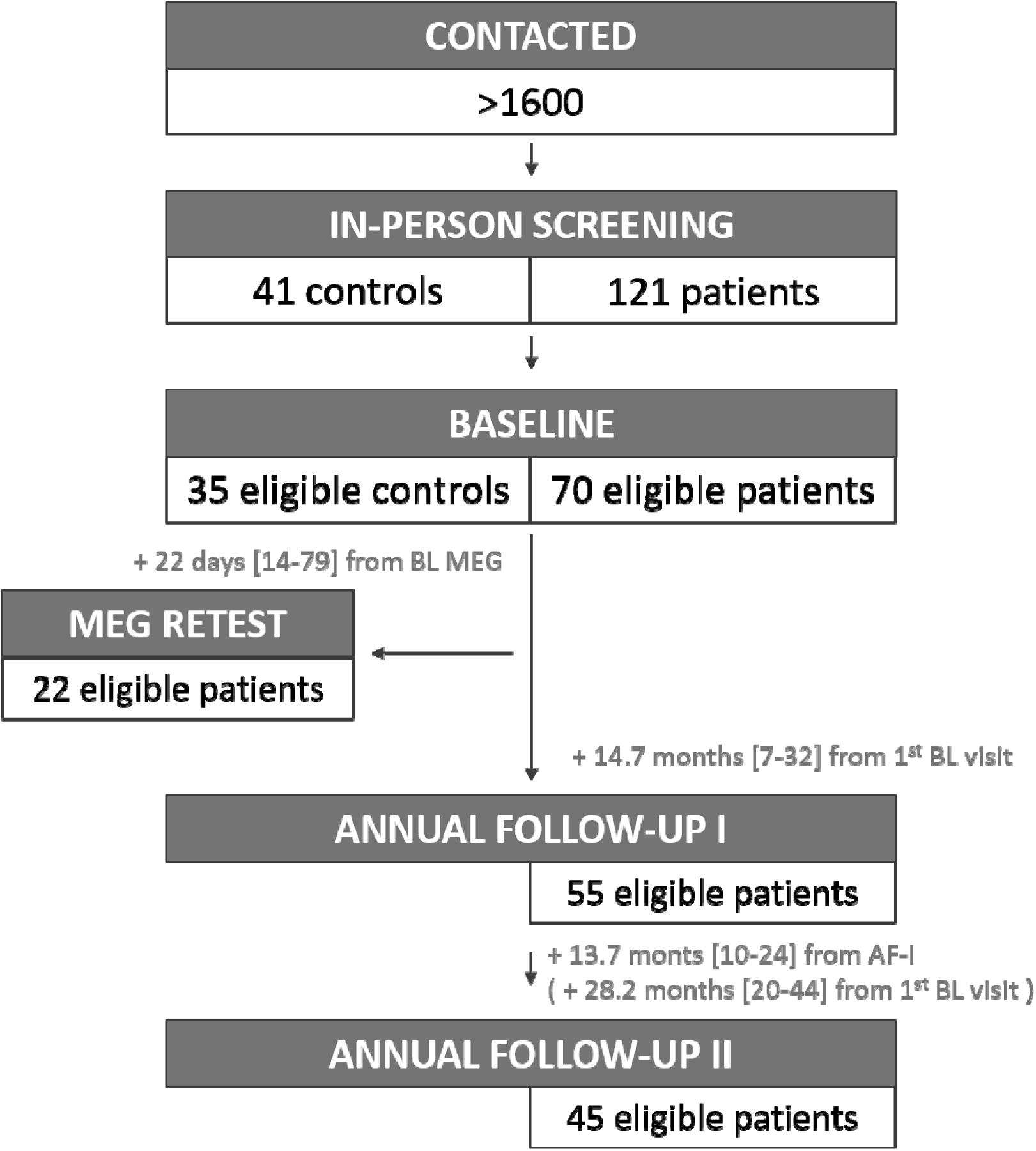
Flow chart of participant recruitment and sample sizes.

Based on these criteria, a total of 70 patients (34 male/36 female) aged 51-84 (mean ± standard (SD) deviation: 73.3 ± 6.9) and 35 healthy controls (21 male/14 female) aged 50-85 (65.4 ± 8.7) were admitted to the main study, of which 69 at Cambridge (49 patients, 20 healthy controls) and 36 at Oxford (21 patients, 15 healthy controls). Distributions of demographics (age at first visit and years of education), biological severity (amyloid status), and cognitive severity are shown in Figure 2. An additional MEG scanner validation dataset (VAL) was acquired at Cambridge, with five healthy participants (3 male/2 female) aged 52-75 (67.8 ± 8.3); see Experimental Design > Simultaneous M/EEG. These are not included in the following analyses.

**Figure 2.**
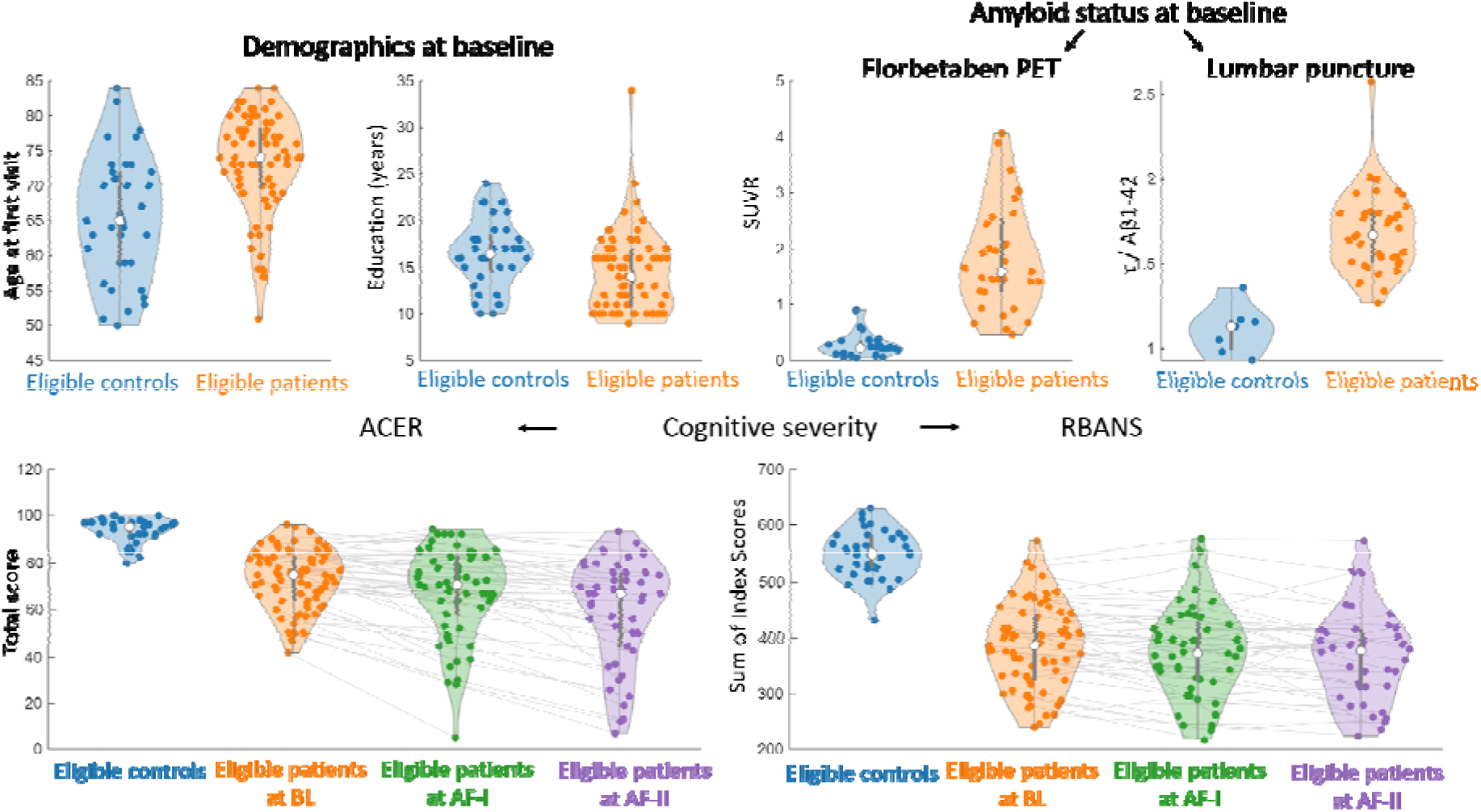
Distributions of demographics and severity of the eligible participants. Healthy controls are shown in blue, and AD/MCI patients are shown in orange (Baseline), green (first annual follow-up), and purple (second follow-up). **Top row left.** Demographics of age at first visit (left panel) and years of education (right). **Top row right.** Amyloid status was determined either by PET or lumbar puncture. Here we show the Standardized Uptake Value Ratio for PET (SUVR; left) and total τ to amyloid β1-42 ratio for CSF (right). **Bottom row.** Shown are the total scores on the Addenbrooke’s Cognitive Examination Revised (ACE-R) and the Sum of Index Scores Repeatable Battery for the Assessment of Neuropsychological Status. Scores are shown for healthy controls at baseline and for patients across all three time points.

### Experimental Design

The experimental design followed the protocol paper (Lanskey et al., 2022), with the caveat of smaller sample size and longer collection timeline mainly related to the Covid-19 pandemic. The protocol prescribed 100 patients and 30 healthy controls at baseline, compared to 70 and 35 included in the dataset, respectively. Participants completed screening and baseline (BL) assessments, biomarker status assessment, clinical and neuropsychological assessments, blood collection, magnetic resonance imaging (MRI), and simultaneous magneto-and electro-encephalography (M/EEG) assessments. Participants in the AD/MCI group were required to have a diagnosis prior to testing and subsequently underwent screening to confirm they met the study-specific inclusion criteria (e.g., biomarker status). For part of the experimental cohort (23 clinical, 3 controls), positive biomarker status had been established previously or simultaneously, with amyloid PET or CSF in clinical or research assessments (their median window between amyloid test and NTAD clinical screening: 8 months, ranging 0-54 months). For the remaining participants, amyloid status was determined during the screening phase using amyloid PET scan (38 patients, 8 healthy controls), or CSF examination (39 patients, 26 healthy controls), depending on the participant’s preference and eligibility. Fifty-five patients completed additional longitudinal assessments, compared to 80 as planned in the protocol paper. Clinical and neuropsychological assessments were repeated twice more at 12–16-month intervals (annual follow-up 1 and 2; AF-I, AF-II). This deviated from the 12-month interval described in the protocol (Lanskey et al., 2022). Covid-related lockdowns and health concerns led to postponed follow-up for many participants as well as higher than expected attrition - the latter increasing statistical power to detect longitudinal changes.

Blood collection, MRI and M/EEG assessments were repeated at the AF-I visit (but not AF-II). AF-I was completed by 55 patients, on average 14.7 months after baseline (range 7-32), and AF-II by 45 patients, on average 28.2 months after baseline (20-44). Participants were only invited for AF-II if they completed AF-I. Additionally, 22 patients completed an early follow-up M/EEG assessment, on average at 22 days after baseline (14-79 days). Lastly, a subset of participants (24 patients, 12 controls) was invited for a UCB-J PET scan. These data were collected under sister studies (MINDMAPS (Mansur et al., 2020) and the SV2A Evaluation in Neurodegeneration and Dementia Research study (SENDER; REC reference 18/EE/0059)) conducted by the same research group rather than under the NTAD protocol, under an aligned protocol, and should be considered a convenience sample (for details, see Data Acquisition > PET).

### Data Acquisition

Figure 3 shows the data completed for each modality (clinical assessment, LP or PET for amyloid status, venepuncture, neuropsychology, M/EEG, MRI, and UCB J PET), separate for each timepoint and group. For each measure, observations were categorised as “collected and eligible”, “not collected during the visit”, “not collected due to ineligibility”, and “collected but excluded from analysis due to not meeting the NTAD inclusion criteria”. The latter category reflects participants whose eligibility status was revised after data collection, resulting in exclusion of all collected data from analysis under current NTAD criteria (e.g., change in diagnosis between sessions or failure to meet scanning eligibility requirements). This is distinct from eligible participants who withdrew (e.g., due to severe disease progression or new health concerns), for whom previously collected eligible data remain analysable. Note that “collected” here indicates that any data were collected during the session, not necessarily that all the data of that session were collected - e.g., if a participant only completed one of the MEG tasks during their visit, they are recorded as “collected”. Further breakdown of data completion is visualised in the modality specific figures (Figure 4 to 8).

**Figure 3.**
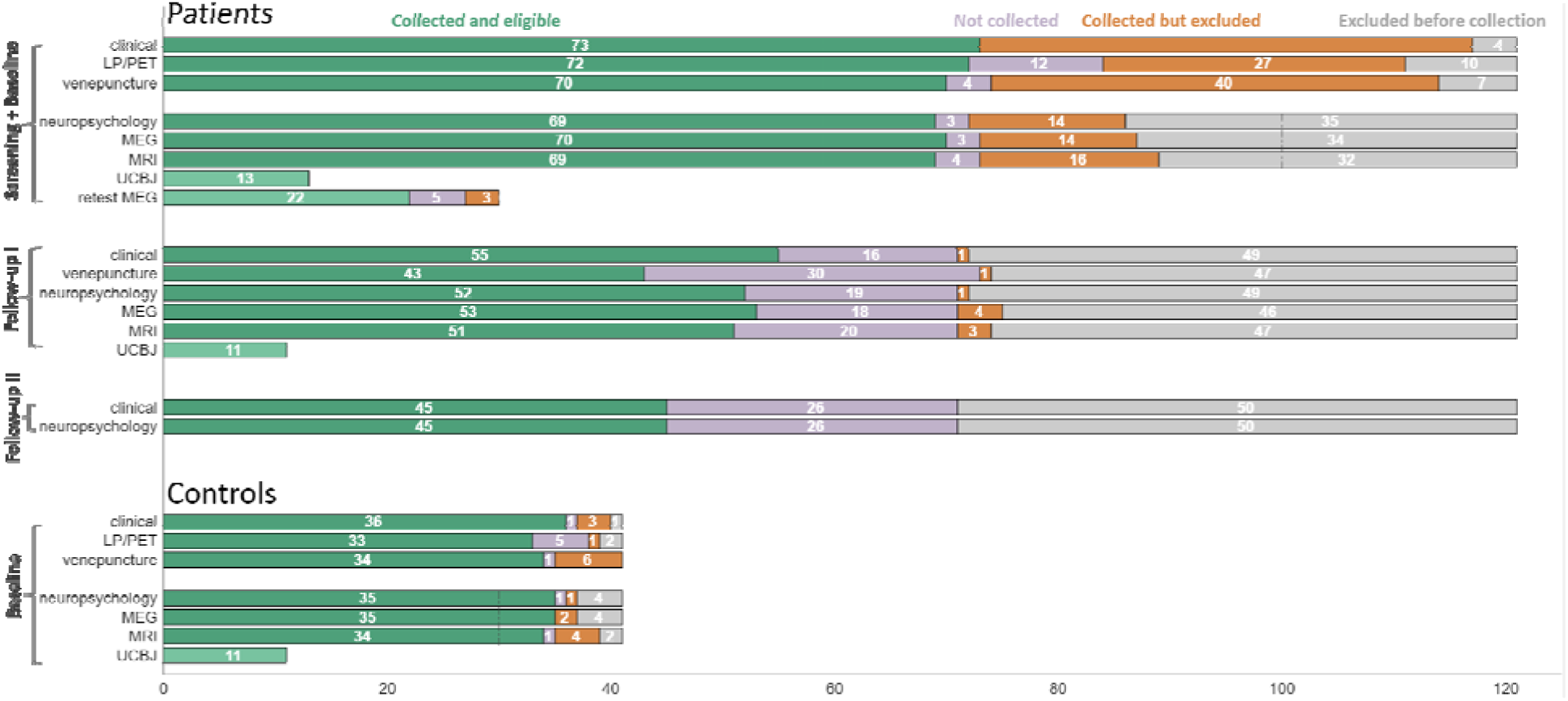
Overview of the data collection for each modality, separate for both groups (patients and controls) and assessment (Screening and Baseline including retest, annual follow-up I, and annual follow-up II). Numbers indicate participant count. Each modality is separated into Collected and eligible (green): data that were collected from eligible participants, Collected but excluded (orange): data that were collected from participants who were deemed ineligible after biomarker testing, Not collected (purple): participants who withdrew or for whom this modality was not successfully collected at all, and Excluded before collection: participants who were excluded based on ineligibility prior to testing. The targeted sample sizes during Baseline from the protocol paper are marked by vertical dashed lines.

**Figure 4.**
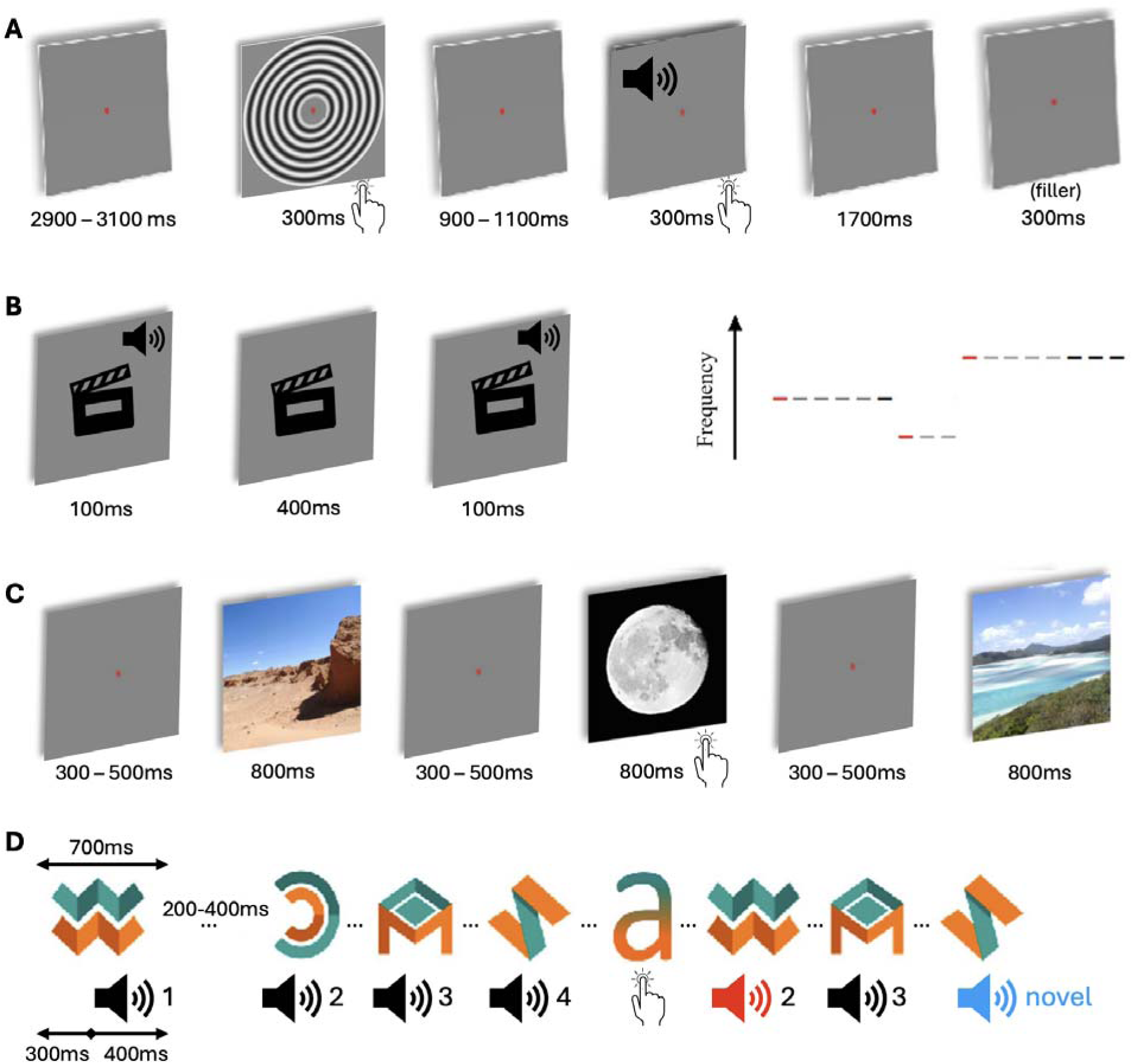
Overview of the four tasks and their trial timings. A) audiovisual task, featuring visual, auditory, and filler trials, B) passive auditory mismatch negativity task, which participants completed while watching a documentary, C) scene repetition task, featuring new and repeated scenes, and D) audiovisual oddball task, featuring learnt pairs, associative deviants, and novel deviants following a training phase.

### M/EEG

MEG data were collected at the Cambridge MRC Cognition and Brain Science Unit (CBU) and Oxford Centre for Human Brain Activity (OHBA). At Cambridge, MEG data were initially collected with a 306-channel Elekta Neuromag VectorView system (15 controls and 47 patients at baseline, 9 patients at AF-I), replaced by a 306-channel MEGIN Triux Neo system (5 controls and 2 patients at baseline, 27 patients at AF-I). At Oxford, MEG data were collected with a Triux Neo system throughout the study. Before the M/EEG assessment, a MEG-compatible passive 64-channel EEG was fitted to the scalp, together with 5 head position indicator (HPI) coils. The location of three fiducials (bridge of the nose, left and right preauricular points), HPI coils, and EEG sensors with respect to the participants’ scalp surface were digitised with a Polhemus OptiTrak system. M/EEG data were collected at a 1000-Hz sampling rate. EEG was not recorded for 17 scans in total (controls at BL: 5, controls at BL-2: 1, patients at BL: 5, patients at TW: 1, patients at AF-I: 5), due to participants not tolerating the cap, head size, or time constraints. At both sites, visual stimuli were presented using a PROPixx projection system (VPixx) situated outside of the magnetically shielded room (MSR), and backprojected onto a semi-translucent screen in front of the participant. Auditory stimuli were presented through silicon tubes from outside the MSR, and fibre-optic button boxes were used to for task responses (see below). In Oxford, gaze and pupil size were recorded using an EyeLink 1000 eye-tracking device (SR Research).

The M/EEG session included four tasks (collected in a consistent order): a simple audio-visual perception task, an auditory mismatch-negativity task, a visual scene repetition task, and a cross-modal audiovisual oddball task (Lanskey *et al*., 2022) (see Table 1 for event triggers and timing). The tasks were followed by recordings of a five-minute period of wakeful rest with eyes open (i.e., “resting state”) and another five-minute period of rest with eyes closed. Eye gaze and pupillometry were only collected during resting state recordings (with eyes open) in Oxford.

**Table 1.**
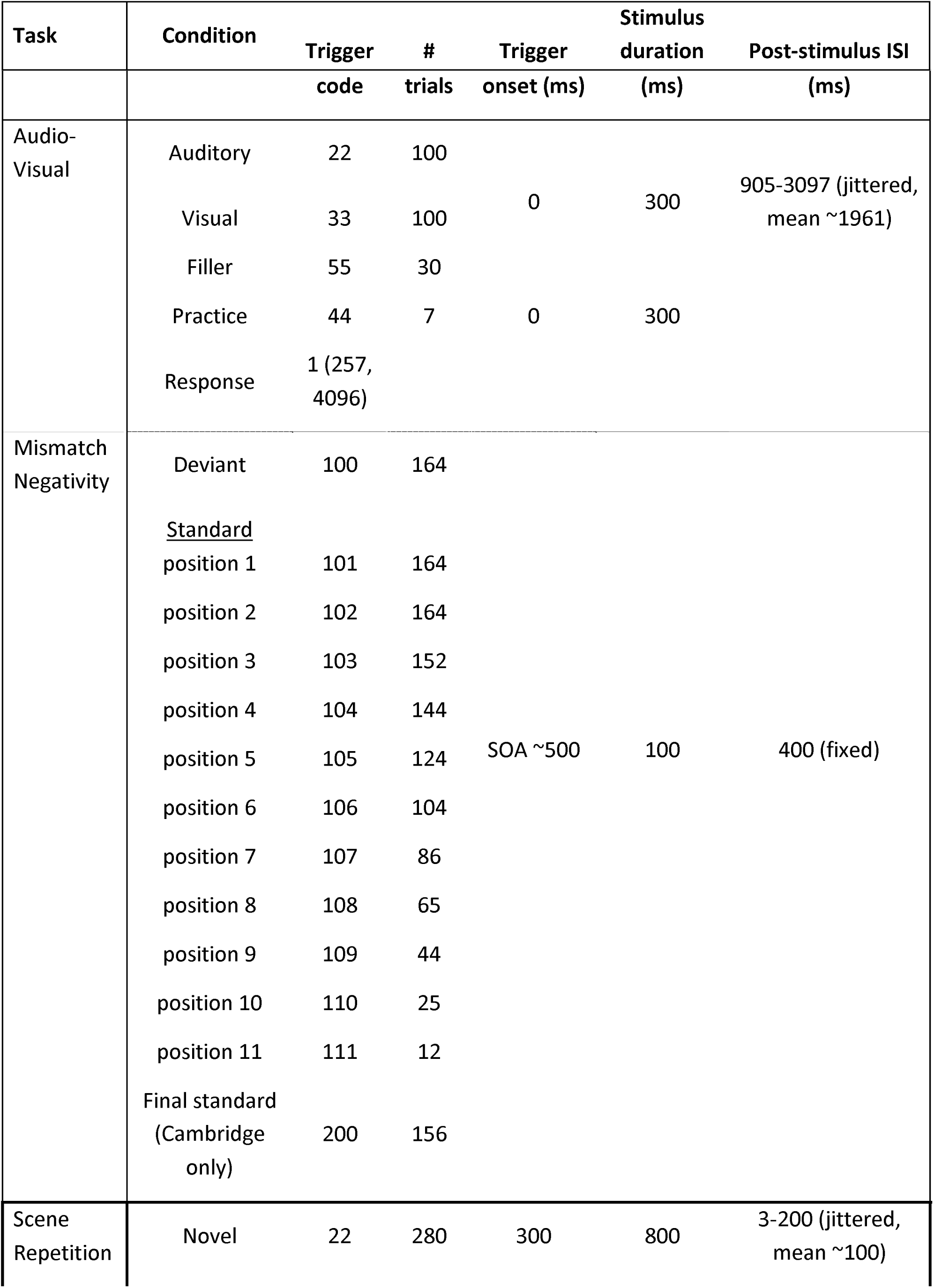
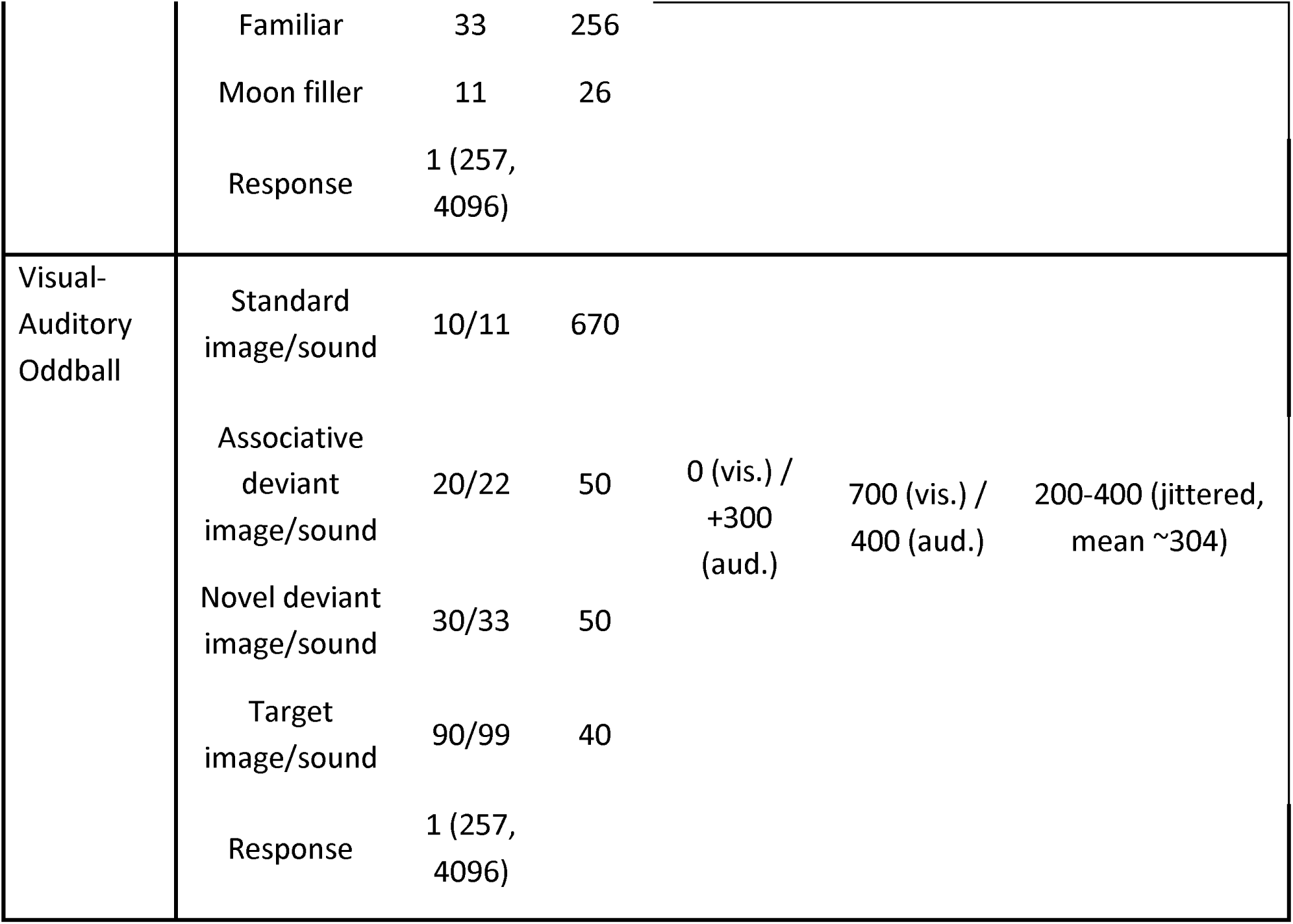
Event durations.

| Task | Condition | Trigger | # | Trigger | Stimulus duration | Post-stimulus ISI |
|---|---|---|---|---|---|---|
|  |  | code | trials | onset (ms) | (ms) | (ms) |
| Audio-Visual | Auditory | 22 | 100 | 0 | 300 | 905-3097 (jittered, mean ~1961) |
|  | Visual | 33 | 100 |  |  |  |
|  | Filler | 55 | 30 |  |  |  |
|  | Practice | 44 | 7 | 0 | 300 |  |
|  | Response | 1 (257, 4096) |  |  |  |  |
| Mismatch Negativity | Deviant | 100 | 164 | SOA ~500 | 100 | 400 (fixed) |
|  | Standard |  |  |  |  |  |
|  | position 1 | 101 | 164 |  |  |  |
|  | position 2 | 102 | 164 |  |  |  |
|  | position 3 | 103 | 152 |  |  |  |
|  | position 4 | 104 | 144 |  |  |  |
|  | position 5 | 105 | 124 |  |  |  |
|  | position 6 | 106 | 104 |  |  |  |
|  | position 7 | 107 | 86 |  |  |  |
|  | position 8 | 108 | 65 |  |  |  |
|  | position 9 | 109 | 44 |  |  |  |
|  | position 10 | 110 | 25 |  |  |  |
|  | position 11 | 111 | 12 |  |  |  |
|  | Final standard (Cambridge only) | 200 | 156 |  |  |  |
| Scene Repetition | Novel | 22 | 280 | 300 | 800 | 3-200 (jittered, mean ~100) |
|  | Familiar | 33 | 256 |  |  |  |
|  | Moon filler | 11 | 26 |  |  |  |
|  | Response | 1 (257, 4096) |  |  |  |  |
| Visual-Auditory Oddball | Standard image/sound | 10/11 | 670 | 0 (vis.) / +300 (aud.) | 700 (vis.) / 400 (aud.) | 200-400 (jittered, mean ~304) |
|  | Associative deviant image/sound | 20/22 | 50 |  |  |  |
|  | Novel deviant image/sound | 30/33 | 50 |  |  |  |
|  | Target image/sound | 90/99 | 40 |  |  |  |
|  | Response | 1 (257, 4096) |  |  |  |  |

Completion rates for each task are shown in Figure 4. Overall completion rates were high for most tasks, indicating the scanning protocol was tolerable for patients even at follow-up. The audiovisual oddball task was the longest task and was more often omitted when experimenters were running out of time. In a few cases, both the mismatch-negativity and the audiovisual tasks were unavailable due to technical issues.

Because of a MEG scanner change at Cambridge in the middle of the NTAD data collection, MEG assessments for 25 patients were completed on different scanners for BL and AF-I sessions. To enable the assessment and correction of scanner effects, a separate MEG scanner validation dataset (VAL) was acquired. Five healthy participants had two M/EEG assessments at the old (Elekta Neuromag VectorView) and two assessments at the new MEG scanner (MEGIN Triux Neo), using the same NTAD M/EEG tasks and data collection procedures. All five validation participants completed the four tasks and two resting states at each scan.

## 3T MRI

MRI data were collected at the CBU and OHBA using the same 3T Siemens PRISMA scanner. The order of MRI sequences was: T1 weighted imaging, T2 weighted fluid attenuated inversion recovery (FLAIR), T2 star weighted imaging, T2 weighted imaging with fat saturation, diffusion weighted imaging, quantitative susceptibility mapping (QSM), resting state functional MRI (fMRI) with eyes open, high resolution T2 weighted imaging of hippocampal subfields, and arterial spin labelling (ASL). Further acquisition details and parameters are provided in Koychev et al. (2018) and Lanskey et al. (2022). Seven participants at baseline (BL) and 11 at the first annual follow up (AF-I) did not complete the full MRI session (Figure 5). Images were visually checked by the operating radiographer during acquisition. Seven hippocampal subfield scans, two T1-weighted scans, and two diffusion-weighted scans were repeated due to excessive motion. In these cases, only the second acquisitions are shared.

**Figure 5.**
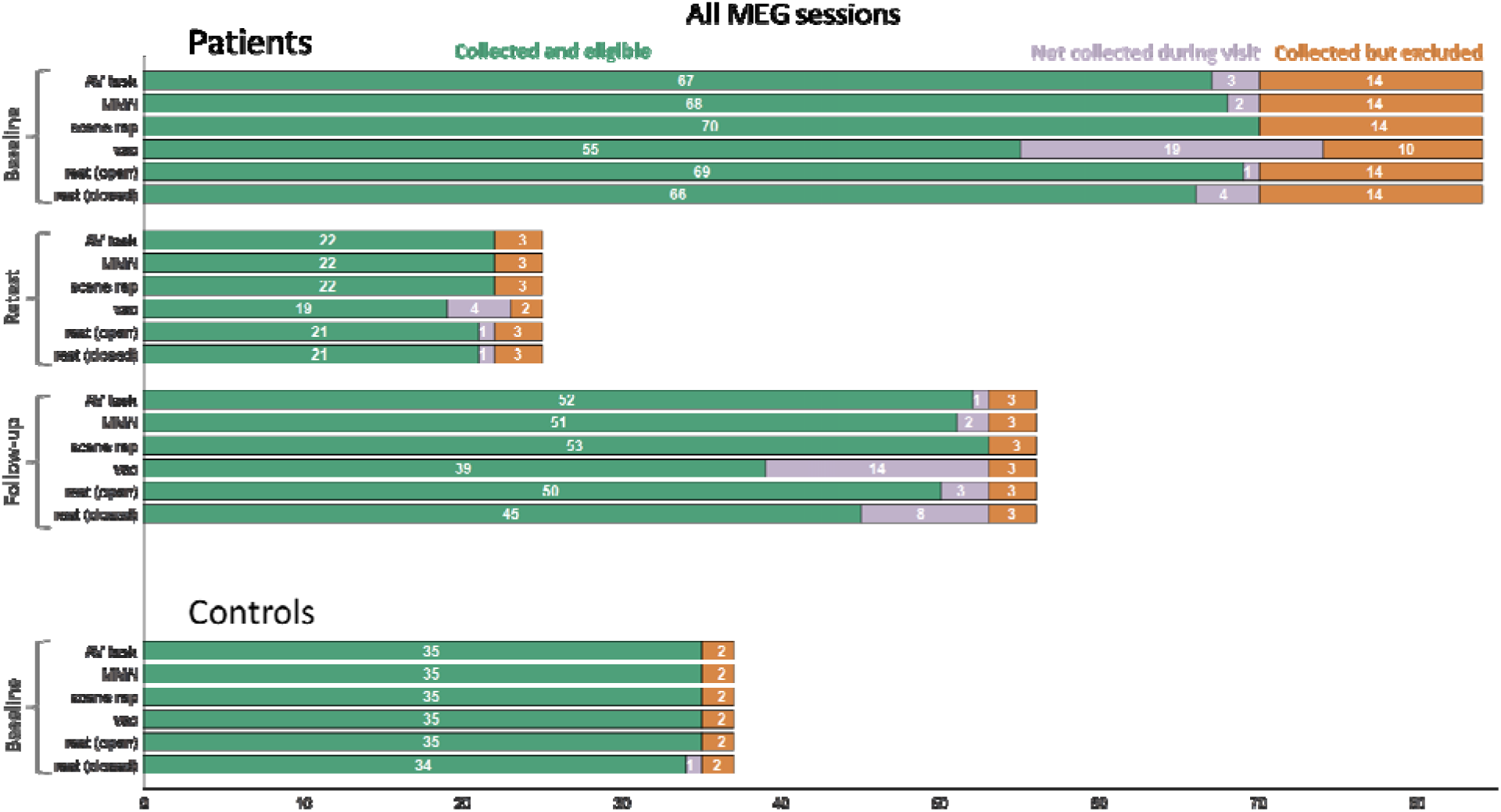
Overview of data completion for the audiovisual perception task (AV task), auditory mismatch negativity (MMN), visual scene repetition (scene rep), and visual-auditory oddball task (VAO) as well as the two resting states (eyes-open and eyes-closed), separate for Baseline (healthy controls and patients), three-week retest and follow-up (patients). Conventions are as in Figure 3.

Two participants initially completed a shorter MRI session (∼15-min) during the screening to obtain structural images required to assess eligibility for a lumbar puncture. One participant subsequently returned at baseline to complete the remaining sequences; the other withdrew before screening was completed. For the participant assessed at both visits, the screening and baseline sequences have been grouped under a single baseline session on XNAT for data sharing. The separate acquisition dates and explanatory comments are retained in the accompanying metadata.

Several inconsistencies in reconstruction availability and acquisition protocol were documented. T1-weighted images were available with and without scanner-based geometric distortion-correction generated from the same acquisition. For two participants, only the distortion-correction reconstruction was available. QSM source data were collected using a single five-echo 3D gradient-echo acquisition. The standard output comprised magnitude and phase images, each reconstructed with and without scanner-based geometric distortion correction. For two participants, only magnitude reconstructions were available, with and without distortion correction; the corresponding phase reconstructions were unavailable. Resting state fMRI comprised a sagittal echo-planar imaging (EPI) acquisition and an associated single-band reference image, together with a short reverse phase-encoding acquisition and corresponding reference image for susceptibility-distortion correction. Three participants missed the reverse phase-encoding acquisition and reference image, while one has these but is missing the EPI acquisition and associated reference image. Eight healthy controls and three patients completed an earlier version of the diffusion protocol, comprising separate acquisitions at nominal b values of 300, 1000 and 2000 s/mm². This was subsequently replaced by a combined multi-shell acquisition at nominal b values of 300, 700 and 2000 s/mm² to align with the revised DFP protocol (Koychev et al., 2018). The earlier protocol is indicated in light green in Figure 5.

### PET

Florbetaben PET scans were collected for 46 participants on a GE Signa PET/MR scanner at the Wolfson Brain Imaging Centre (Cambridge) or a GE D710 PET/CT scanner at the Churchill Hospital (Oxford) to establish amyloid positivity (Rowe et al., 2017; Sabri et al., 2015). For another 9 participant’s Florbetaben SUVR values were obtained from prior studies conducted by the same research group and shared with the NTAD study team.

¹¹C UCB J PET scans (Botermans, 2023; Toyonaga et al., 2019) were collected with a 3T PET/MR at the Cambridge Wolfson Brain Imaging Centre (BL: 2 patients, 3 healthy controls; AF I: 2 patients) or at Invicro London (BL: 5 patients, 8 healthy controls; AF I: 4 patients, 1 healthy control), or with PET/CT at Invicro London (BL: 6 patients; AF I: 5 patients). Some participants completed the scan as part of their baseline assessments (13 patients, 11 controls) and others at the time of their first follow up assessments (11 patients, 1 control). Processed SUVR values across 90 nodes, alongside the raw UCB J data, will be made available in a future release.

### Cerebrospinal Fluid

Lumbar punctures were performed for 66 participants at the Cambridge Clinical Research Centre and Oxford Health NIHR Clinical Research Facility to collect a maximum of 10 ml of clear CSF. For another 16 participants, CSF AD-biomarkers values were obtained from prior studies conducted by the same research group or from local clinics, and shared with the NTAD study team. Total τ, phosphorylated τ, and Aβ1^-40^/Aβ1^-42^ were quantified at the Institute of Neurology, UCL, to determine study eligibility.

### Clinical assessment

Clinical assessments were taken at baseline, AF-I, and AF-II (Figure 7). We share data from the revised Addenbrooke’s cognitive examination (ACE-R; Mioshi et al., 2006) and Clinical Dementia Rating (CDR; Hughes et al., 2018; Morris et al., 1993). Participants also underwent a longer clinical interview at screening and shorter interviews at follow-up sessions to assess medical history and medication usage. We also share data from questionnaires collected for all participants, including healthy controls, comprising self-assessments (Geriatric Depression Scale (GDS; Ishihara et al., 2011) and State-Trait Anxiety Inventory (STAI; Spielberger et al., 1970) and study partner assessments (Amsterdam Instrumental Activities of Daily Living scale (iADL; Jutten et al., 2017) and Mild Behavioural Impairment Checklist (MBI-C; Ismail et al., 2017)). As expected, data completion was high for the ACE-R and CDR and more prominent for the questionnaires. Data completion remained high for the follow-up sessions (Figure 6).

**Figure 6.**
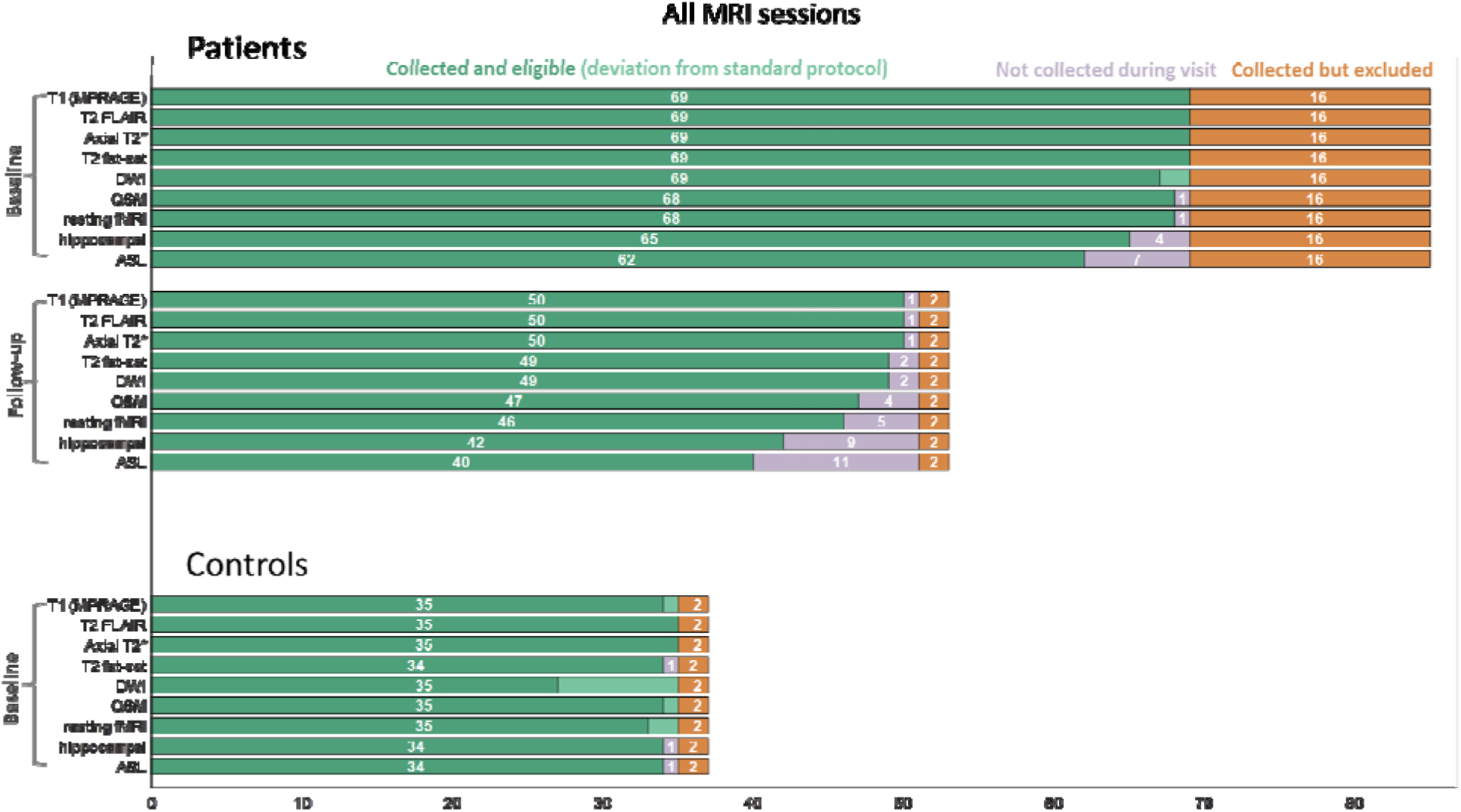
Overview of data completion for T1-weighted (T1 MPRAGE), T2 FLAIR, T2*-weighted, T2-weighted with fat saturation (T2 fat-sat), diffusion-weighted (DWI), quantitative susceptibility mapping (QSM), resting state fMRI, hippocampal subfields, and arterial spin labelling (ASL), separate for baseline (healthy controls and patients), three-week retest and follow-up (patients). Conventions are as above.

**Figure 7.**
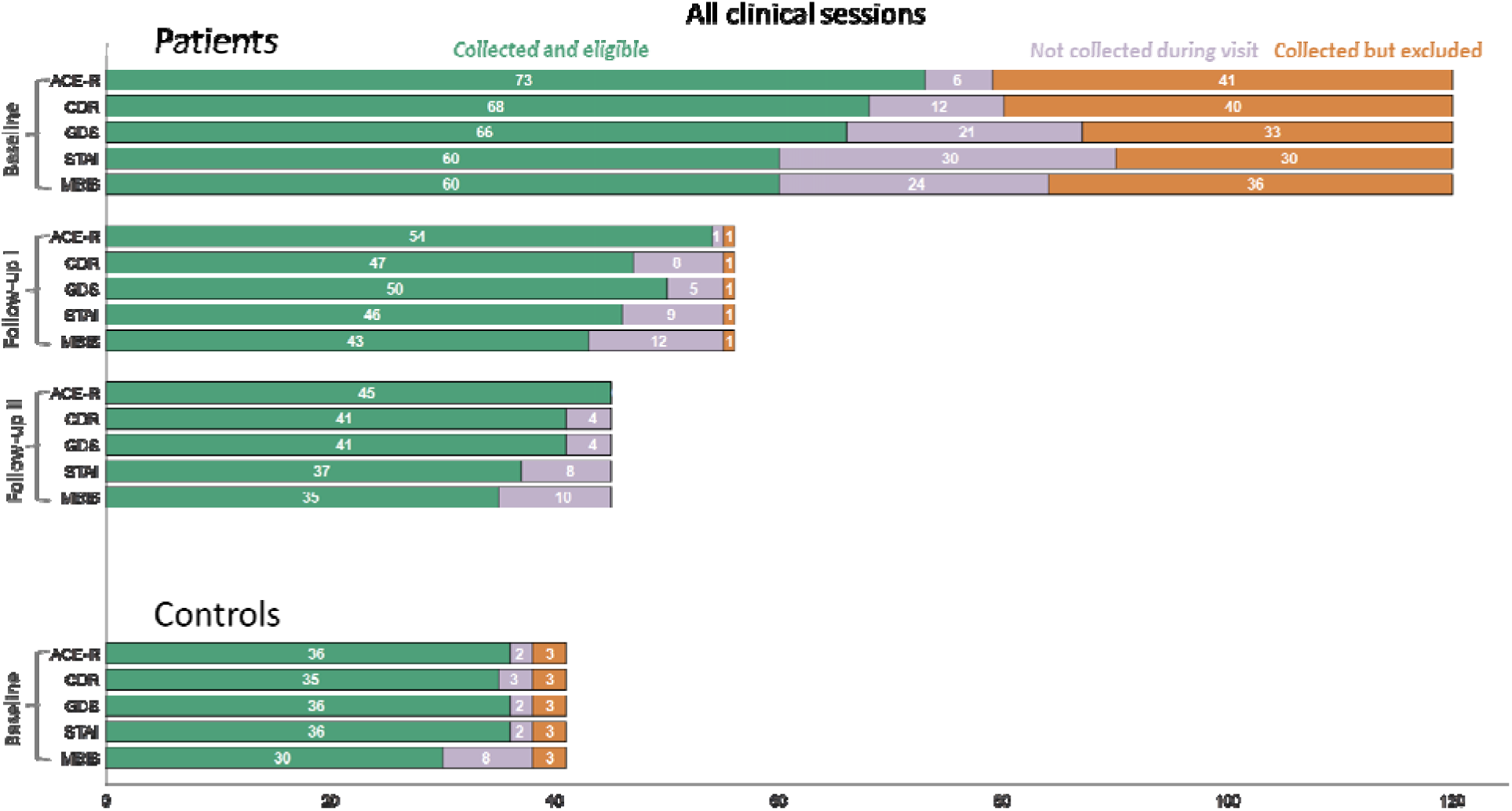
Overview of data completion for clinical assessments and self-and partner-assessed questionnaires. Conventions are as above.

**Figure 8.**
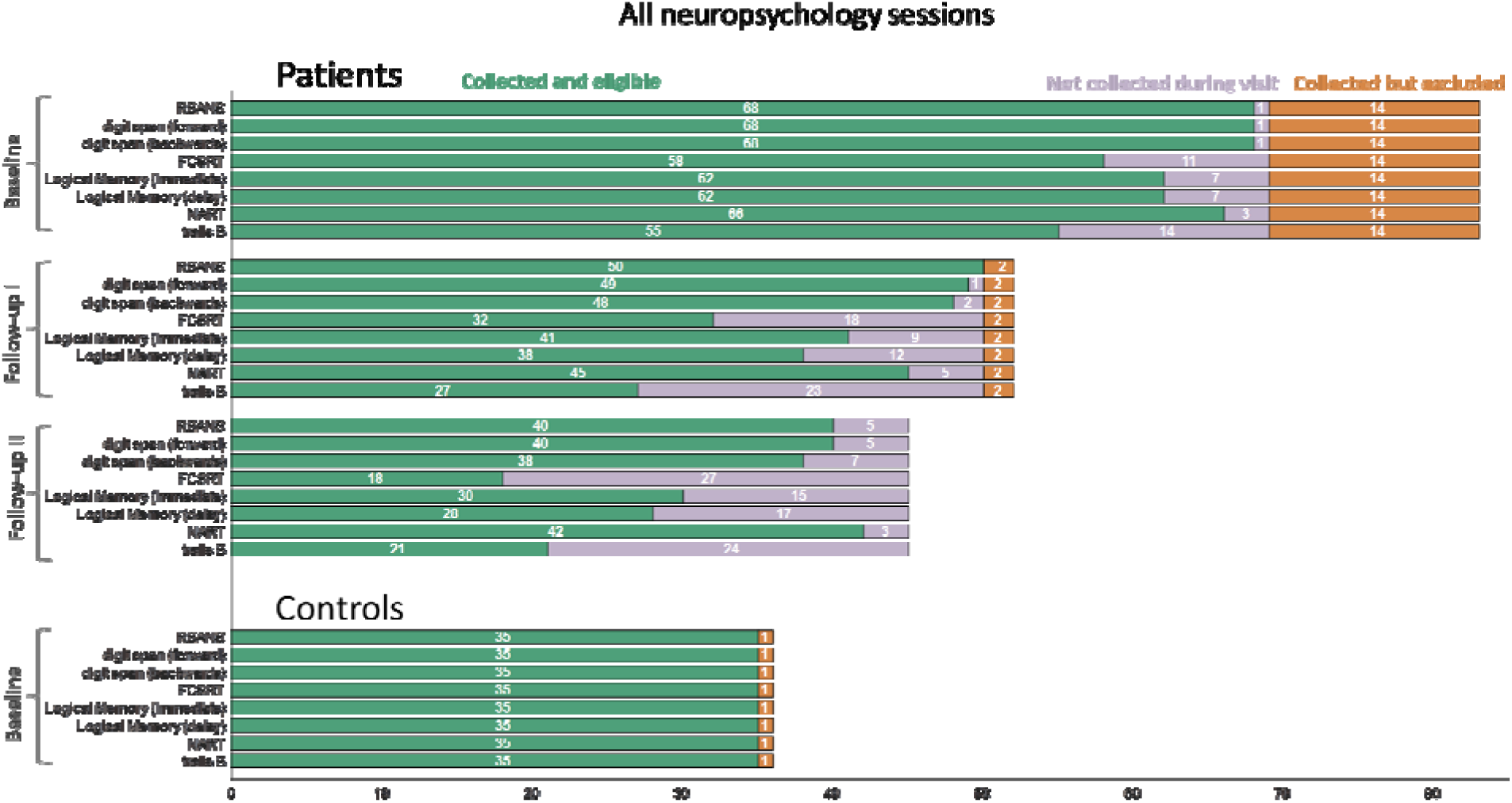
Overview of data completion for the pen-and-paper neuropsychological assessments. Conventions are as above.

Missing values in questionnaire data were addressed through imputation where appropriate. For questionnaires with <25% missing items, missing responses were imputed at the item level separately for each participant. These cases were typically questionnaires completed on paper, where individual items were likely skipped unintentionally. After imputation, the total questionnaire score was calculated across the observed and imputed items. For questionnaires with ≥25% missing items, imputation was not performed due to insufficient information – these cases were recorded as “not collected” in the final dataset. For the GDS (binary items) and STAI (Likert scale), each item contributes equally. We therefore imputed missing items with the person-specific median value of the completed items. This does not work for the iADL and MBI-C as it contains conditional sub-items. For the iADL, the total questionnaire score with data imputation was calculated by taking the mean of completed items, multiplying by the number of missing items, and rounding to the nearest integer. For the MBI-C, imputed scores were calculated for the binary items first and conditional severity scores second. Across all measures, imputed scores were closely aligned with observed data, suggesting minimal impact on between-subject variance while improving statistical power. For transparency, we provide both the variables with and without imputed scores in the shared dataset. Exact details of the data imputations are reported in Table 2.

**Table 2.**

| Scale | # session with missing items | [min-max] # items of missing items (%) | # sessions with imputed items (% missing items) | imputed score |
|---|---|---|---|---|
| GDS | 11<br>(BL: 1 control, 5 patients; AF-I: 4, AF-II: 2) | 1-2 (7-13%) | 11 (7-13%) | Median value |
| iADL | 5<br>(BL: 3 patients; AF-II: 2) | 1-11 (7-37%) | 3 (3-10%) | Mean value multiplied by number of missing items, and rounded to nearest integer |
| MBI-C | 6<br>(BL: 3 patients; AF: 1 ; AF-II: 2) | 3-18 (8-54%) | 2 (8%) | Binary items: modal response<br>Severity scores: median (if other severity scores available) or minimum score |
| STAI trait | 3<br>(BL: 2 patients, 1 control) | 1-3 (5-15%) | 3 (5-15%) | Median value |
| STAI scale | 2<br>(BL: 1 patient; AF-II: 1) | 1-6 (5-30%) | 1 (5%) | Median value |

### Neuropsychological assessment

Neuropsychological assessments were performed at baseline, AF-I, and AF-II timepoints. Experimenters assessed participants using the following paper-and-pen tests: the Repeatable Battery for the Assessment of Neuropsychological Status (RBANS; Randolph et al., 1998), digit span backwards taken from the Wechsler Adult Intelligence Scale (Wechsler, 1981), Free and Cued Selective Reminding Test (FCSRT; Grober et al., 2018), Logical Memory taken from the Wechsler Memory Scale (3rd edition; Wechsler, 1999), National Adult Reading Test (NART; Nelson & Willison, 1991), digit symbol substitution taken from the Wechsler Adult Intelligence Scale (Wechsler, 1981), and Trail Making Test B (Bowie & Harvey, 2006). Participants also completed tasks on a tablet: the Four Mountains Test (4MT; Hartley et al., 2007), as well as simple and five-choice reaction time, paired associates learning, rapid visual processing, and spatial working memory tasks from the Cambridge Neuropsychological Test Automated Battery (CANTAB; Cambridge Cognition, 2019).

Completion rates were high for the first two tasks (RBANS and Digit Span), but lower for the remaining tasks, likely reflecting both increased task difficulty as well as participant fatigue. Particularly, a substantial number of participants could not complete the FCSRT, logical memory, and Trail Making tasks. This pattern was more pronounced at follow-up sessions (Figure 8). Tablet-based tasks were considered most challenging by participants and correspondingly showed the lowest completion rates, with substantially reduced completion rates at follow-up sessions (Figure 9).

**Figure 9.**
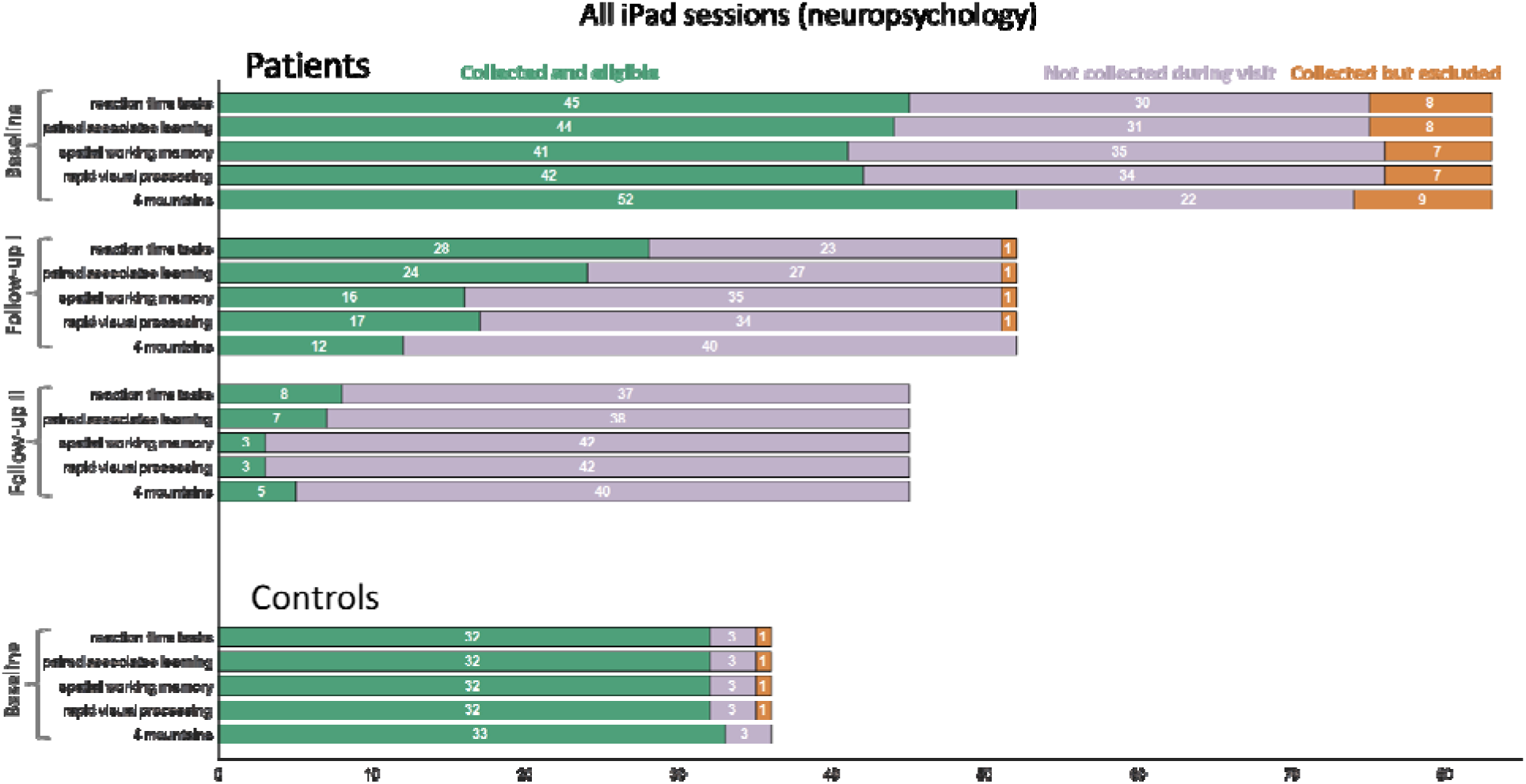
Overview of data completion for the tablet-based neuropsychological assessments. Conventions are as above.

### Venepuncture for blood samples

Blood samples were obtained at BL in the following order: 2.7 mL in sodium citrate tubes (SCT) for plasma, 5 mL in serum separator tubes (SST) for serum, 10 mL in EDTA tubes for DNA extraction, a further 10 mL in EDTA tubes for plasma and buffy coat processing, and two 2.5 mL PAXgene tubes for RNA preservation. At AF-I, blood drawing was repeated for SCT, SST, and whole blood tubes for all participants, and for DNA and RNA for a subset. In Oxford, safety bloods were also collected at BL to assess eligibility as per local requirements, with the results reviewed and confirmed by the local doctor and principal investigator (PI). The blood tests included a coagulation screen, vitamin B12, ferritin, urea and electrolytes (U&Es), liver function tests (LFTs), thyroid function tests (TSTs), and full blood count (FBC). At the Cambridge site, research blood samples were processed and stored by the local biorepository. At the Oxford site, safety bloods were processed locally and the results shared to determine eligibility. Research bloods were processed accordingly, stored in freezers, and shipped on dry ice to the biorepository in Cambridge.

Blood collection was not successful in all cases (e.g., due to poor venous access). In three cases, follow-up bloods could successfully be drawn at AF-II but not AF-I. Blood processing also occasionally failed. We therefore report the tubes successfully processed and held by the local biorepository in Cambridge (Figure 10). Any failures are reported as “not collected during visit”. Initial analyses will focus on proteomics, with Alamar NULISAseq neurodegeneration (CNS120) and inflammation (250) panels.

**Figure 10.**
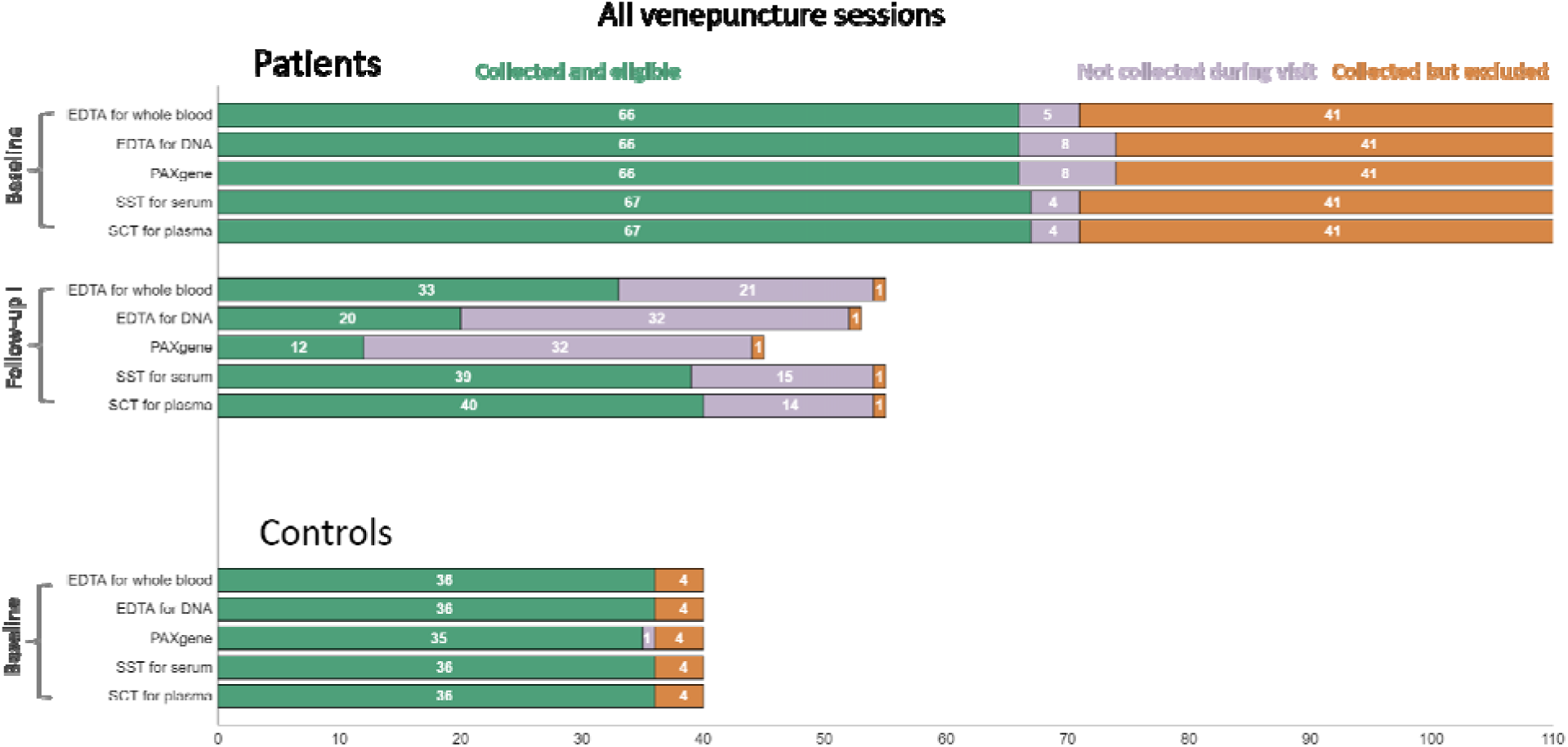
Overview of data completion for the venepuncture for blood samples. Conventions are as above.

### Data Records

[To be supplemented with details of DPUK pipelines, including defacing]

https://github.com/OHBA-analysis/NTAD_data_paper2026

Imaging data are organised using the Brain Imaging Data Structure (BIDS). Data are organised hierarchically by participant, session, and task/sequence. Additional data including those arising from neuropsychological and cognitive assessments, results from blood and biomarker analysis, and metadata are included as CSV files accompanying metadata/data dictionaries in JSON format. These extra files are further explained in the README file of the GitHub repository (https://github.com/OHBA-analysis/NTAD_data_paper2026). MRI and PET data are stored in NifTI format with accompanying metadata stored in JSON format.

### MEG Preprocessing

MEG data are stored in Functional Imaging File (FIF) format at various levels of preprocessing: raw (i.e., before any preprocessing – together with EEG data), and after applying various stages of MaxFilter and further preprocessing (see below).

MEGIN’s MaxFilter was applied to the MEG data in three steps (https://imaging.mrc-cbu.cam.ac.uk/meg/maxpreproc): first, bad channels were identified using the *-autobad* option. Second, Signal Space Separation with temporal extension (tsss; Taulu & Simola, 2006) was applied with movement compensation depending on the availability of valid head position indicator (HPI) measurements, or without movement compensation otherwise. Third, the data were projected to a common coordinate frame, by estimating the head origin from the head shape points, and moving this to [0, 13,-6] relative to the device origin, as advised by the MaxFilter manual (Elekta Oy, 2010).

MEG data were further preprocessed after the second stage of MaxFilter processing, see Figure 11. We used OSL-Ephys (Van Es et al., 2025; Quinn et al., 2022) v0.10.0 and MNE-Python (Gramfort et al., 2013; 2014) v1.3.1 for preprocessing the data, as follows (https://github.com/OHBA-analysis/NTAD_data_paper2026). Note that the DPUK-supplied data after the third MaxFilter stage (projection to a common coordinate frame) have not been preprocessed further, nor have the EEG data.

**Figure 11.**
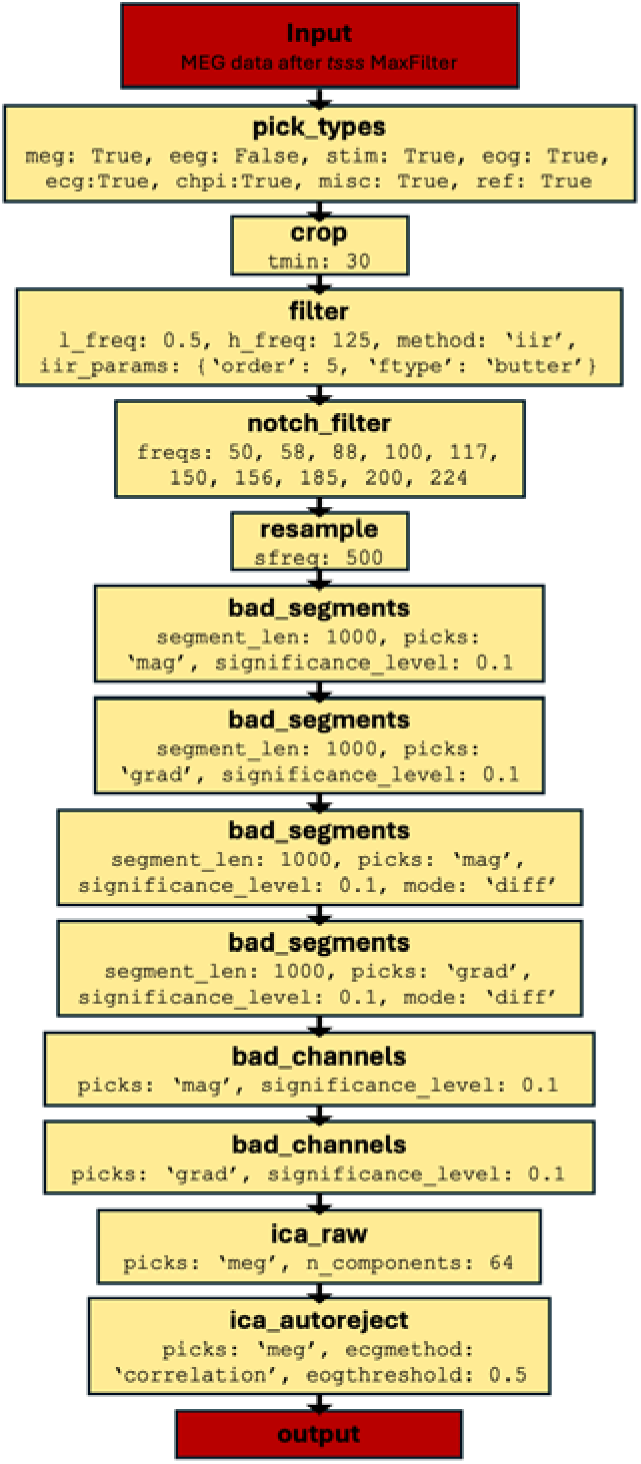
Overview of the MEG preprocessing pipeline, applied after the MaxFilter tsss step.

The first 30 s of MEG data – during which the HPI signals were initialised – were discarded. The data were band-pass filtered between 0.5 and 250 Hz using a 5-order Butterworth infinite impulse-response filter. Line noise at 50 Hz and its harmonics were removed with a notch filter (2 Hz width), and additionally, interference from unknown sources was notched at [58, 88, 117, 156, 185, 224] Hz. Following down-sampling to 500 Hz, periods of high variance were detected using a generalized extreme studentized deviate (gESD) test (Posner, 1983) on 2-s windows of the raw signal, and 2-s windows of the temporal derivative (significance level 0.1). Bad channels were also detected using a gESD test with the same settings (separately for magnetometers and gradiometers). Finally, cardiac and ocular artefacts were removed using temporal independent component analysis (ICA). ICA was fitted to a copy of the data that was high-pass filtered at 1 Hz, using all good MEG channels (each sensor type standardised by its standard deviation) and excluding segments marked as bad. These data were reduced to their first 64 principal components, and FastICA was used to estimate 64 independent components. Components were flagged as ocular if their time course correlated with the EOG (|r| > 0.5), and as cardiac if their correlation with the ECG was an outlier relative to the other components (z > 3). All flagged components were checked visually. Only the rejected components were removed from the original (0.5 Hz high-passed) data; principal components beyond the first 64 were kept, so the PCA step did not reduce the rank of the data.

### Technical Validation

The MEG preprocessing outcomes are summarised in Table 3. Overall, MaxFilter with movement compensation was successful for 1025/1060 files. The other 35 were rerun without movement compensation. Furthermore, an average of 7.49% (standard deviation (SD): 6.40%; range: 0 to 68.50%) of data segments were marked as bad because of high variance artifacts, and on average 2.11 gradiometers (SD: 3.89; range: 0 to 21) and 1.13 magnetometers (SD: 2.23; range: 0 to 11) were marked as bad, and 3.06 ICs (SD: 1.14; range: 0 to 10) were removed. Event types, timings, numbers, and trigger codes are summarised in Table 1. Delays between trigger onset and visual stimulus onset were 12.0 and 10.0 ms in Cambridge and Oxford, respectively; delays between trigger onset and auditory stimulus onset were 64.0 and 68.5 ms, respectively. These delays were measured by comparing the trigger channel to the corresponding recorded photodiode and audio signal channels on empty room pilot data (see GitHub). These estimates may be less precise for scans recorded in Cambridge after June 2020 for visual stimuli and December 2022 for auditory stimuli due to small changes in hardware. We note that different estimates have been used in a prior publication (Lanskey et al., 2025); see the Data documentation for more details. Forty-four scans collected at Cambridge after the scanner change had incorrectly stored trigger values, but these can be recovered by restructuring the event array (see Data Documentation).

**Table 3.** Summary metrics of the MEG preprocessing pipeline, indicating the percentage of the data marked as “bad” (i.e., containing high variance or related to signals not of interest). Every column shows the mean (standard deviation) [range] across files. Grad.: gradiometers; Mag.: magnetometers; IC: independent components.

| **Task** | **# Files** | **Segments (%)** | **Grad. (#)** | **Mag. (#)** | **IC (#)** |
|---|---|---|---|---|---|
| All Tasks | 1060 | 7.49 (6.40)<br>[0-68.50] | 2.11 (3.89)<br>[0-21] | 1.13 (2.23)<br>[0-11] | 3.06 (1.14)<br>[0-10] |
| Audio-Visual | 183 | 6.87 (5.67)<br>[0-32.78] | 2.21 (3.93)<br>[0-19] | 1.11 (2.20)<br>[0-11] | 3.10 (0.97)<br>[0-6] |
| Mismatch Negativity | 181 | 7.46 (5.07)<br>[0-27.24] | 2.14 (3.56)<br>[0-16] | 1.29 (2.29)<br>[0-11] | 3.03 (1.13)<br>[0-7] |
| Scene repetition | 186 | 7.55 (6.97)<br>[0.13-68.50] | 2.40 (4.57)<br>[0-21] | 1.18 (2.43)<br>[0-11] | 2.82 (1.09)<br>[0-7] |
| Visual-Auditory Oddball | 155 | 7.47 (5.84)<br>[0.73-43.88] | 2.11 (3.81)<br>[0-19] | 1.13 (2.18)<br>[0-11] | 2.92 (1.11)<br>[0-5] |
| Resting state<br>Eyes Open | 182 | 8.11 (6.85)<br>[0-30.98] | 2.03 (3.65)<br>[0-21] | 1.29 (2.22)<br>[0-11] | 3.01 (1.00)<br>[1-6] |
| Eyes Closed | 173 | 7.50 (7.65)<br>[0-38.67] | 1.76 (3.69)<br>[0-17] | 0.76 (2.01)<br>[0-11] | 3.46 (1.40)<br>[1-10] |

To provide a descriptive validation that the MEG recordings captured the sensory and cognitive responses targeted by the experimental protocols, we examined global field power (GFP) during the four task paradigms and power spectral density (PSD) during the resting state recordings. These analyses were based on all MEG data (i.e., all sessions across all NTAD participants) and were intended as technical validation of the recorded data rather than as inferential tests of experimental, diagnostic group, or longitudinal effects.

For all analyses, the MEG data were resampled to 500 Hz. For the evoked response analyses, gradiometer data were low pass filtered at 30 Hz and epoched around events of interest. Epochs were averaged across trials and baseline corrected using the 200 ms period preceding stimulus onset. GFP was calculated as the root mean square across all gradiometers and baseline corrected at the session level before averaging within participants and subsequently across participants. For the resting state recordings, PSDs were estimated using Welch’s method with a Hamming window and 4-s segments. Session level spectra were averaged across gradiometers, converted to decibels, and subsequently averaged within participants and across participants.

The resulting group averaged responses showed characteristic stimulus locked MEG activity across all four task paradigms (Fig. 12a to d). The audiovisual task produced clear responses following both auditory and visual stimulation, while the mismatch negativity paradigm showed differentiated GFP time courses for standard and deviant tones. The scene repetition and visual auditory oddball paradigms likewise produced clear stimulus locked responses, with condition dependent differences visible in the group averaged time courses. These analyses demonstrate that the task recordings contain measurable evoked responses and condition related signal variation suitable for subsequent analyses.

**Figure 12.**
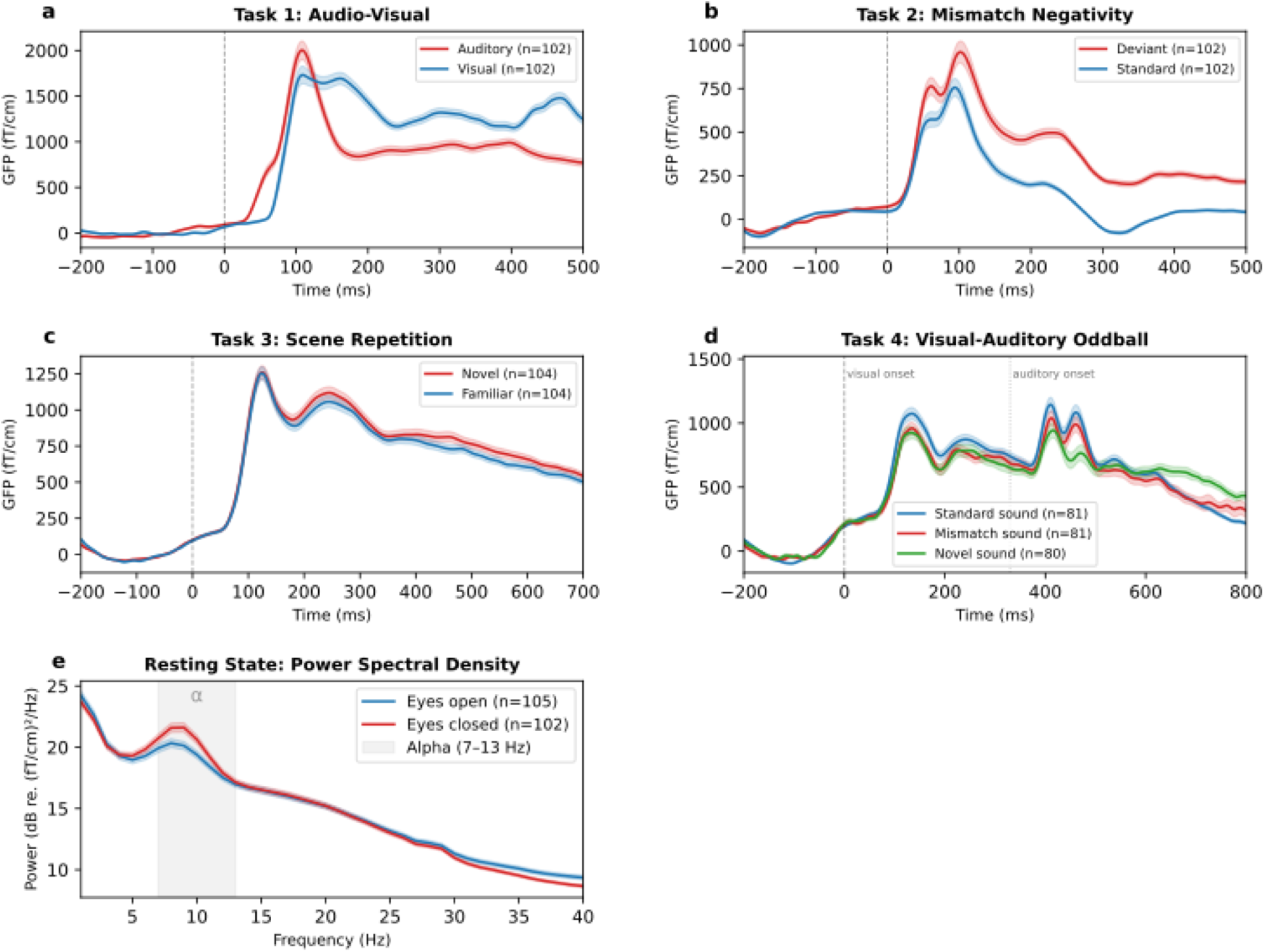
Technical validation of task evoked responses and resting state spectral characteristics. a to d, Group averaged global field power (GFP) calculated across gradiometers for the audiovisual task (a), mismatch negativity task (b), scene repetition task (c), and visual auditory oddball task (d). In (d), conditions correspond to standard trials, trials containing an auditory deviant, and trials containing a novel deviant stimulus. Vertical dotted lines indicate stimulus onset; in (d), the later auditory stimulus onset is additionally indicated. (e) Group averaged power spectral density (PSD) during eyes open and eyes closed resting state recordings. PSDs (Welch, 4 s windows) were averaged over all gradiometers and converted to dB per recording, then averaged across visits within each participant. The shaded region denotes the canonical alpha frequency range, 7 to 13 Hz. Solid lines indicate the participant mean and shaded regions indicate the standard error of the mean. Values of n in the legends indicate the number of participants contributing data to each condition.

Resting state spectra showed the expected 1/f broadband decrease in power with increasing frequency, along with a prominent alpha band peak between 7 and 13 Hz (Figure 12e). Alpha band power was descriptively greater during eyes closed than eyes open recordings, consistent with a well-established feature of resting human MEG. Together, these analyses provide basic validation that the dataset captures canonical evoked and spontaneous electrophysiological features targeted by the acquisition protocol.

### Scanner change at the Cambridge site

In August 2020, the Cambridge Vectorview was replaced by a Triux Neo (43 NTAD sessions acquired after the change). To assess its impact, five healthy volunteers completed the full protocol twice on each system, with sessions on the same scanner 0–15 days apart and the two scanners ∼10 months apart. We correlated mean-centred per-channel features between all recordings: log-power spectra for resting state and evoked responses for the tasks. On every measure, recordings from the same participant were far more similar than those from different participants, on either scanner (Figure 13). Similarity across scanners was slightly lower than on the same scanner (gradiometers, mean r = 0.34–0.67 vs 0.49–0.74), though this cannot be disentangled from the longer interval between scans on different scanners (∼10 months vs ≤15 days); the scanner effect is therefore at most this small. The participant could be identified across scanners in 111 of 117 recordings (95%; chance 20%). We recommend including scanner as a covariate when combining Cambridge data from before and after the change.

**Figure 13.**
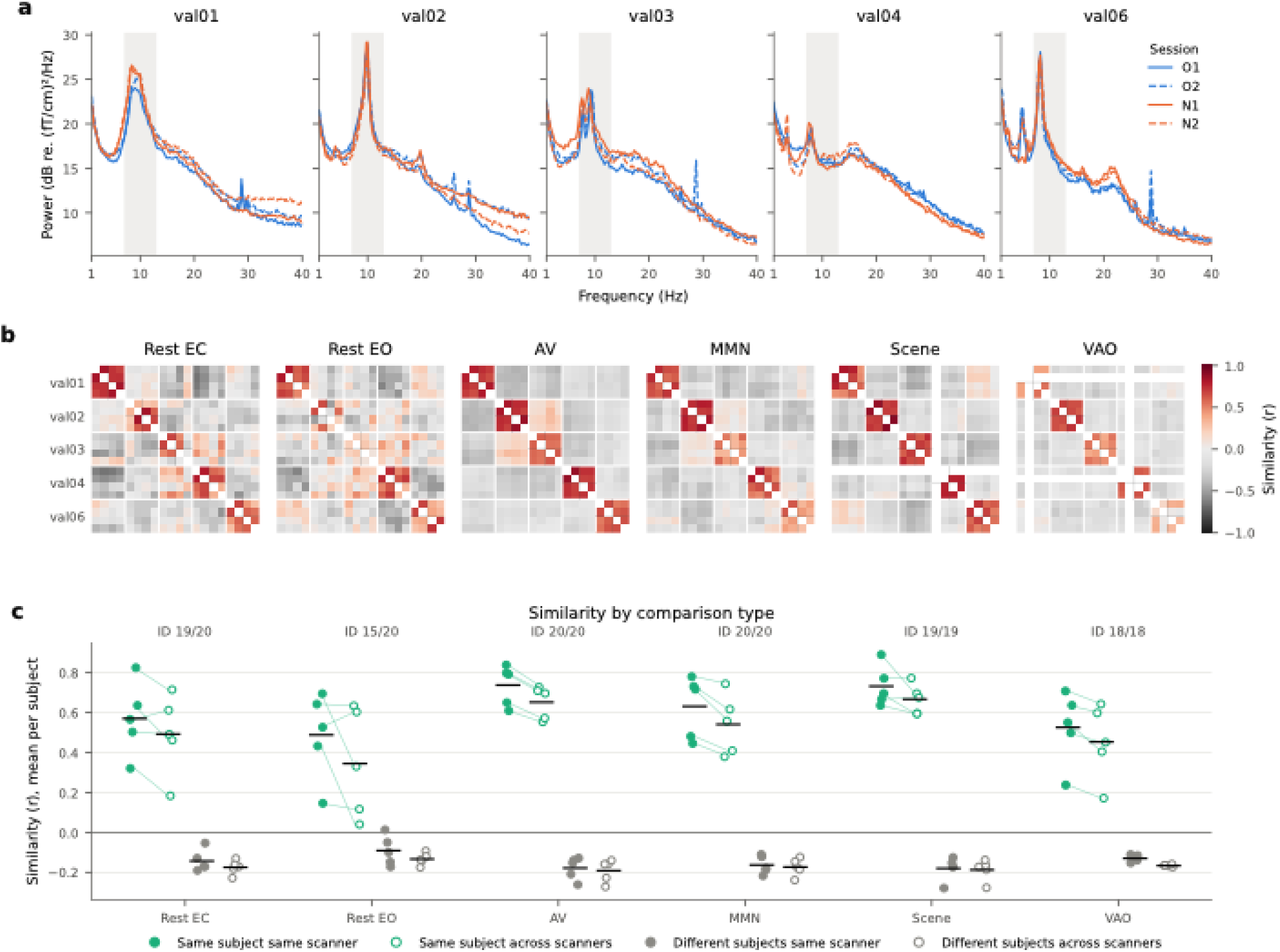
Cambridge scanner change validation. Five healthy volunteers were scanned twice on the Vectorview (O1, O2) and twice on the Triux Neo (N1, N2), with sessions on the same scanner 0–15 days apart and the two scanners ∼10 months apart. **a**, Eyes-closed resting spectra for each participant (gradiometers, sensor average). **b**, Similarity (Pearson r) between all 20 recordings for each measure, ordered by participant, then session; white marks the diagonal and recordings that were not acquired. **c**, Mean similarity per participant for each comparison type; numbers above each measure give cross-scanner identification accuracy: the proportion of recordings whose most similar recording from the other scanner came from the same participant (chance = 20%). Rest EC/EO, eyes closed/open; AV, audio-visual; MMN, mismatch negativity; Scene, scene repetition; VAO, visual–auditory oddball.

### Usage Notes

The current data release comprises nine CSV files containing participant demographics, study metadata, and clinical and cognitive assessments. Descriptions of variables are provided in the accompanying *NTAD Data Documentation.* This document also includes detailed instructions on how to use the data. We have shared data irrespective of participant suitability for the planned NTAD analyses. Users can apply different criteria appropriate to their own research aims. A variable indicating inclusion in NTAD analyses is provided in the participant overview and can be used to restrict analyses to participants meeting these criteria.

To summarise, the spreadsheet *“NTAD_metadata_all”* provides a comprehensive overview of all consented participants and all scheduled study sessions. It specifies the data completion as summarised in Figures 4 and A1-A4: sessions attended by each participant, sessions that were acquired but deemed unusable, including reasons for exclusion, and sessions not attended, with reasons where available. An additional comments field provides further detail on participant ineligibility or withdrawal. Further participant-level details are available in the accompanying CSV files. Together, these files provide a complete record of each participant.

We provide a GitHub repository with additional data, scripts, and usage notes, which may be helpful when analysing the data. For example, MaxFilter calibration files are provided for each scanner to enable the user to MaxFilter the MEG data with alternative settings. We also provide measurements of the auditory and visual delays with respect to the trigger timing (separately for each site). Furthermore, we share functions to recover incorrectly recorded triggers and to align triggers across sites.

To correctly read the event information from the MEG trigger channel, we recommend the following usage (in MNE-Python): mne.find_events(‘STI101’, mask=4352 (256|4096), mask_type=’not_and’, uint_cast=True).

## Data Availability

Data are findable at https://portal.dementiasplatform.uk/. They can be compared or aligned with other studies using the Cohort Explorer, https://portal.dementiasplatform.uk/data/cohort-explorer/. Applications to access the data are made via https://data.dpuk.ukserp.ac.uk/Apply. At time of publication, average approval time is three weeks.

## Code Availability

Custom scripts used for MEG data preprocessing, quality assessment and technical validation, together with supporting documentation, are publicly available through GitHub at https://github.com/OHBA-analysis/NTAD_data_paper2026. The GitHub page also contains links to related NTAD publications (and accompanying code repositories) published by the consortium.

## Author Contributions

Conceptualization: JBR, ACN, MWW, RNH, SL, MT.

Data curation: MVE, MNP, AJQ, JP, AK, TT, EK, JL, MK, CA, MN.

Formal analysis: MVE, MNP.

Funding acquisition: JBR, ACN, MWW, KDS.

Investigation: MVE, MNP.

Methodology: MVE, MNP, AJQ, AK, EK, JL, HF, VR, ACN, MWW, JBR.

Project administration: MVE, MNP, AJQ, JP, AK, TT, EK, JL, MK, CA, HF, VR, ACN, MWW, JBR.

Resources: ACN, MWW, JBR, SL, MT.

Software: MVE, MNP, AJQ.

Supervision: MVE, MNP, AJQ, HF, ACN, MWW, JBR.

Validation: MVE, MNP, CA.

Visualization: MVE, MNP.

Writing, original draft: MVE, MNP.

Writing, review and editing: MVE, MNP, AJQ, JP, AK, TT, EK, JL, MK, CA, HF, VR, KDS, RNH, ACN, MWW, JBR.

## Acknowledgements

MVE and MNP have contributed equally to the current work and have the right to list their name first when referencing the work. We are very grateful to the participants and their family members for their contributions to the study. We also would like to thank the DPUK Data Team, particularly Chloe Tulip, Cat Morris, and Emma Squires, for their support in data sharing. We thank the following people for their help during data collection: Sven Braeutigam, Anna Camera, Lara Bolte, Oscar Haines, Juliet Semple, Jon Campbell, Nicky Aikin, David Parker, Seb Rieger, Marius Mada, Clare Cook, Lucy Browns, Tina Emery, Lucy McGregor, Steve Elridge, the CCRC Team, and the nurses and medical staff.

## Ethics statement

The New Therapeutics in Alzheimer’s Disease (NTAD) study had ethical approval from the Cambridge Central Research Ethics Committee, East of England (REC reference 18/EE/0042).

## Appendix A: Glossary

| 4MT | Four Mountains Test (completed on iPad) |
|---|---|
| ACER | Addenbrooke’s Cognitive Examination – Final Revised (2005) |
| AD | Alzheimer’s Disease |
| AF-I | Annual Follow-up I |
| AF-II | Annual Follow-up II |
| Aβ | Amyloid-β protein |
| ASL | Arterial spin labelling; MRI sequence |
| AV task | Audiovisual task during M/EEG recording |
| bioFIND | Multi-site resting state magnetoencephalography dataset including participants with mild cognitive impairment and healthy controls |
| BIDS | Brain Imaging Data Structure, standardised format for neuroimaging datasets |
| BL | Baseline |
| Cam-CAN | Cambridge Centre for Ageing and Neuroscience study, containing data along the adult lifespan and 12-year follow-up data |
| CANTAB | Cambridge Neuropsychological Test Automated Battery, recorded on iPad during neuropsychology session |
| CDR | Clinical Dementia Rating Scale |
| CSF | Cerebrospinal fluid |
| Digit span backwards | Digit span backwards task taken from the Wechsler Adult Intelligence Scale, assessed by experimenter during neuropsychology session on pen-and-paper |
| Digit span substitution | Digit symbol substitution as taken from the Wechsler Adult Intelligence Scale, assessed by experimenter during neuropsychology session on pen-and-paper |
| DFP | Deep and Frequent Phenotyping, a sister study from NTAD |
| DPUK | Dementia Platform United Kingdom |
| DWI | Diffusion-weighted imaging; MRI sequence |
| EDTA | Ethylenediaminetetraacetic Acid tube for blood collection. Used in NTAD to collect DNA as well as plasma and buffy coat. |
| EEG | Electroencephalography |
| FCSRT | Free and Cued Selective Reminding Test, containing 4x4 items which participants are asked to recall in a free recall and cued recall phase |
| GDS | Geriatric Depression Scale, self-assessed questionnaire during clinical sessions with pen-and-paper or iPad |
| hippocampal | MRI sequence designed to acquire a high resolution image of the hippocampus |
| iADL | Amsterdam instrumental activities of daily living questionnaire, assessed by study partner during clinical sessions with pen-and-paper or iPad |
| Logical Memory | Logical memory as taken from the Wechsler memory scale, consisting of an immediate and delayed (+20 min) phase |
| LP | Lumbar puncture |
| MBI-C | Mild Behavioral Impairment Checklist, assessed by study partner during clinical sessions with pen-and-paper or iPad |
| MCI | Mild cognitive impairment |
| MEG | Magnetoencephalography |
| MMN | Roving mismatch negativity task during M/EEG recording |
| MMSE | Mini–mental state examination |
| MRI | Magnetic Resonance Imaging |
| NART | National Adult Reading Test, assessed by experimenter during neuropsychology session on pen-and-paper |
| OMEGA | Open MEG Archive (OMEGA) |
| PAL | Paired Associates Learning task from CANTAB. Participants completed trials with 2, 4, 6, and 8 patterns. |
| PAXgene | Blood tube type used to collect RNA |
| PET (PET/MRI, PET/CT) | Positron emission tomography to measure tracer uptake, combined either with MR or computed tomography (PET/CT) for coregistration to a brain image |
| PREVENT-AD | Pre-symptomatic Evaluation of Experimental or Novel Treatments for Alzheimer's Disease dataset containing longitudinal MEG data on at-risk participants |
| QSM | Quantitative susceptibility mapping; MRI sequence |
| RBANS | Repeatable Battery for the Assessment of Neuropsychological Status, assessed by experimenter during neuropsychology session on pen-and-paper |
| rest | Resting state during M/EEG recording (including both eyes-open and eyes-closed) |
| Resting fMRI | Resting state functional MRI acquisition. |
| RVP | Rapid Visual Information Processing task from CANTAB |
| RT | Reaction Time task from CANTAB. Participants performed both the simple RT task (1 target) and 5-choice RT task. |
| scene rep | Scene repetition task during M/EEG recording |
| SCT | Sodium citrate tube for blood collection (used to collect plasma) |
| SST | Serum separator tube for blood collection (used to collect serum) |
| STAI | State Trait Anxiety Inventory comprising separate trait and state scales, self-assessed questionnaire during clinical sessions with pen-and-paper or iPad |
| SUVR | Standardized Uptake Value Ratio; standard PET measure assessing uptake of a radiotracer in a target compared to a reference region |
| SWM | Spatial Working Memory task from CANTAB. Participants completed trials with 4, 6, and 8 boxes. |
| T1 (MPRAGE) | T1-weighted Magnetization-Prepared Rapid Gradient Echo; MRI sequence |
| T2 FLAIR | T2-weighted Fluid-Attenuated Inversion Recovery; MRI sequence |
| T2* | Axial T2 star weighted; MRI sequence |
| T2 fat-sat | T2-weighted with fat saturation; MRI sequence |
| Trials-B | Trail Making Test Part B, assessed during neuropsychology session on pen-and-paper |
| TW | Three-week retest MEG scan (planned to be 2 weeks in protocol, but 3 weeks on average in practice) |
| UCB-J | 11C-UCB-J PET tracer to measure synaptic vesicle glycoprotein 2A (SV2A) density |
| VAL | MEG scanner validation dataset acquired at Cambridge during the scanner replacement |
| vao | visual-auditory oddball task during M/EEG recording |
| WAND | Welsh Advanced Neuroimaging Database, multi-modal neuroimaging database along the adult lifespan |

## References

Adams, N.E., Hughes, L.E., Phillips, H.N., Shaw, A.D., Murley, A.G., Nesbitt, D., Cope, T.E., Bevan-Jones, R., Passmonti, L. & Rowe, J.B. (2020). GABA-ergic dynamics in human frontotemporal networks confirmed by pharmaco-magnetoencephalography. Journal of Neuroscience, 40(8), 1640–1649. 10.1523/JNEUROSCI.1689-19.2019

Bauermeister, S., Orton, C., Thompson, S., Barker, R. A., Bauermeister, J. R., Ben-Shlomo, Y., Brayne, C., Burn, D., Campbell, A., Calvin, C., Chandran, S., Chaturvedi, N., Chêne, G., Chessell, I. P., Corbett, A., Davis, D. H. J., Denis, M., Dufouil, C., Elliott, P., Fox, N., Hill, D., Hofer, S. M., Hu, M. T., Jindra, C., Kee, F., Kim, C. H., Kim, C., Kivimäki, M., Koychev, I., Lawson, R. A., Linden, G. J., Lyons, R. A., Mackay, C., Matthews, P. M., McGuiness, B., Middleton, L., Moody, C., Moore, K., Na, D. L., O’Brien, J. T., Ourselin, S., Paranjothy, S., Park, K. S., Porteous, D. J., Richards, M., Ritchie, C. W., Rohrer, J. D., Rossor, M. N., Rowe, J. B., Scahill, R., Schnier, C., Schott, J. M., Seo, S. W., South, M., Steptoe, A., Tabrizi, S. J., Tales, A., Tillin, T., Timpson, N. J., Toga, A. W., Visser, P. J., Wade-Martins, R., Wilkinson, T., Williams, J., Wong, A., & Gallacher, J. E. J. (2020). The Dementias Platform UK (DPUK) Data Portal. European Journal of Epidemiology, 35(6), 601–611. 10.1007/s10654-020-00633-4

Botermans, W., Koole, M., Van Laere, K., Savidge, J. R., Kemp, J. A., Sunaert, S., Duffy, M. M., Ramael, S., Cesura, A. M., D’Ostilio, K., Gossen, D., Madsen, T. M., Lodeweyckx, T., & de Hoon, J. (2023). SDI-118, a novel procognitive SV2A modulator: First-in-human randomized controlled trial including PET/fMRI assessment of target engagement. Frontiers in Pharmacology, 13, Article 1066447. 10.3389/fphar.2022.1066447

Bowie, C. R., & Harvey, P. D. (2006). Administration and interpretation of the Trail Making Test. Nature Protocols, 1(5), 2277–2281. 10.1038/nprot.2006.390

Busche, M. A., & Hyman, B. T. (2020). Synergy between amyloid-β and tau in Alzheimer’s disease. Nature Neuroscience, 23(10), 1183–1193. 10.1038/s41593-020-0687-6

Cacabelos, R. (2022). What have we learnt from past failures in Alzheimer’s disease drug discovery? Expert Opinion on Drug Discovery, 17(4), 309–323. 10.1080/17460441.2022.2031784

Carson RE, Naganawa M, Toyonaga T, Koohsari S, Yang Y, Chen MK, Matuskey D, Finnema SJ. Imaging of Synaptic Density in Neurodegenerative Disorders. J Nucl Med. 2022 Jun;63(Suppl 1):60S–67S. PMID: 35649655; PMCID: PMC12530506. 10.2967/jnumed.121.263201

Cope, T. E., Hughes, L. E., Phillips, H. N., Adams, N. E., Jafarian, A., Nesbitt, D., Assem, M., Woolgar, A., Duncan, J., & Rowe, J. B. (2022). Causal evidence for the multiple demand network in change detection: auditory mismatch magnetoencephalography across focal neurodegenerative diseases. Journal of Neuroscience, 42(15), 3197–3215. 10.1523/JNEUROSCI.1622-21.2022

Cambridge Cognition. *CANTAB® [Cognitive assessment software]*, 2019.

Demetriou, I., Attaheri, A., Bingham, T., Duckett, W., Bridge, L., Raykov, P., Tsvetanov, K.A., Correia, M., Apšvalka, D., Crespo-Garcia, M., Campbell, K., Morcom, A., Mitchell, D.J., Rowe, J., Wolpe, N., Henderson, S.E., Cam-CAN & Henson R. (2025). The Cambridge Centre for Ageing and Neuroscience (Cam-CAN) longitudinal study protocol: Phase 4 (“Enrichment”) and Phase 5 (“Rescan”). MedrXiv. d 10.1101/2025.05.06.25327023

DeKosky, S. T., & Scheff, S. W. (1990). Synapse loss in frontal cortex biopsies in Alzheimer’s disease: Correlation with cognitive severity. Annals of Neurology, 27(5), 457–464. 10.1002/ana.410270502

Elekta Oy. (2010). *MaxFilter™ user’s guide* (Version 2.2).

Van Es, M. W. J., Gohil, C., Quinn, A. J., & Woolrich, M. W. (2025). osl-ephys: A Python toolbox for the analysis of electrophysiology data. Frontiers in Neuroscience, 19, Article 1522675. 10.3389/fnins.2025.1522675

Gramfort, A., Luessi, M., Larson, E., Engemann, D. A., Strohmeier, D., Brodbeck, C., Parkkonen, L., & Hämäläinen, M. S. (2013). MEG and EEG data analysis with MNE-Python. Frontiers in Neuroscience, 7, Article 267. 10.3389/fnins.2013.00267

Gramfort, A., Luessi, M., Larson, E., Engemann, D. A., Strohmeier, D., Brodbeck, C., Goj, R., Jas, M., Brooks, T. L., Parkkonen, L., & Hämäläinen, M. S. (2014). MNE software for processing MEG and EEG data. NeuroImage, 86, 446–460. 10.1016/j.neuroimage.2013.10.027

Gorgolewski, K. J., Auer, T., Calhoun, V. D., Craddock, R. C., Das, S., Duff, E. P., Flandin, G., Ghosh, S. S., Glatard, T., Halchenko, Y. O., Handwerker, D. A., Hanke, M., Keator, D., Li, X., Michael, Z., Maumet, C., Nichols, B. N., Nichols, T. E., Pellman, J.,…Poldrack, R.A. (2016). The brain imaging data structure, a format for organizing and describing outputs of neuroimaging experiments. Scientific Data, 3, Article 160044. 10.1038/sdata.2016.44

Grober, E., Veroff, A. E., & Lipton, R. B. (2018). Temporal unfolding of declining episodic memory on the Free and Cued Selective Reminding Test in the predementia phase of Alzheimer’s disease: Implications for clinical trials. *Alzheimer’s & Dementia: Diagnosis*, Assessment & Disease Monitoring, 10, 161–171. 10.1016/j.dadm.2017.12.004

Hartley, T., Bird, C. M., Chan, D., Cipolotti, L., Husain, M., Vargha-Khadem, F., & Burgess, N. (2007). The hippocampus is required for short-term topographical memory in humans. Hippocampus, 17(1), 34–38. 10.1002/hipo.20240

Hughes, L. E., Adams, N. E., Rouse, M. A., Naessens, M., Shaw, A., Murley, A. G., Cope, T. E., Holland, N., Nesbitt, D., Street, D., Whiteside, D. J., & Rowe, J. B. (2025). GABAergic modulation of beta power enhances motor adaptation in frontotemporal lobar degeneration. Alzheimer’s & Dementia, 21(5), e14531. 10.1002/alz.14531

Hughes, C. P., Berg, L., Danziger, W. L., Coben, L. A., & Martin, R. L. (1982). A new clinical scale for the staging of dementia. The British Journal of Psychiatry, 140(6), 566–572. 10.1192/bjp.140.6.566

Hughes, L.E., Henson, R.N., Pereda, E., Bruña, R., López-Sanz, D., Quinn, A.J., Woolrich, M.W., Nobre, A.C., Rowe, J.B., Maestú, F. & BioFIND Working Group (2019). Biomagnetic biomarkers for dementia: A pilot multicentre study with a recommended methodological framework for magnetoencephalography. *Alzheimer’s & Dementia: Diagnosis*, Assessment & Disease Monitoring, 11, 450–462. 10.1016/j.dadm.2019.04.009

Hughes, L. E., Rittman, T., Regenthal, R., Robbins, T. W., & Rowe, J. B. (2015). Improving response inhibition systems in frontotemporal dementia with citalopram. Brain, 138(7), 1961–1975.

Ishihara, T., & Terada, S. (2011).*[Geriatric Depression Scale (GDS)]*. *Nihon Rinsho, 69*(Suppl. 8), 455–458.

Ismail, Z., Agüera-Ortiz, L., Brodaty, H., Cieslak, A., Cummings, J., Fischer, C. E., Gauthier, S., Geda, Y. E., Herrmann, N., Kanji, J., Lanctôt, K. L., Miller, D. S., Mortby, M. E., Onyike, C. U., Rosenberg, P. B., Smith, E. E., Smith, G. S., Sultzer, D. L., Lyketsos, C., & NPS Professional Interest Area of the International Society to Advance Alzheimer’s Research and Treatment. (2017). The Mild Behavioral Impairment Checklist (MBI-C): A rating scale for neuropsychiatric symptoms in pre-dementia populations. Journal of Alzheimer’s Disease, 56(3), 929–938. 10.3233/JAD-160979 [pmc.ncbi.nlm.nih.gov], [alzforum.org]

Ittner, L. M., Ke, Y. D., Delerue, F., Bi, M., Gladbach, A., van Eersel, J., Wölfing, H., Chieng, B. C., Christie, M. J., Napier, I. A., Eckert, A., Staufenbiel, M., Hardeman, E., & Götz, J. (2010). Dendritic function of tau mediates amyloid-β toxicity in Alzheimer’s disease mouse models. Cell, 142(3), 387–397. 10.1016/j.cell.2010.06.036

Jafarian, A., Assem, M. K., Kocagoncu, E., Lanskey, J. H., Fye, H., Williams, R., Quinn, A. J., Pitt, J., Raymont, V., Lowe, S., Singh, K. D., Woolrich, M., Nobre, A. C., Henson, R. N., Friston, K. J., & Rowe, J. B. (2025). Neurophysiological progression in Alzheimer’s disease: Insights from dynamic causal modelling of longitudinal magnetoencephalography. Human Brain Mapping, 46(8), e70234. 10.1002/hbm.70234

Jafarian, A., Assem, M. K., Kocagoncu, E., Lanskey, J. H., Williams, R., Cheng, Y.-J., Quinn, A. J., Pitt, J., Raymont, V., Lowe, S., Singh, K. D., Woolrich, M., Nobre, A. C., Henson, R. N., Friston, K. J., & Rowe, J. B. (2024). Reliability of dynamic causal modelling of resting-state magnetoencephalography. Human Brain Mapping, 45(10), e26782. 10.1002/hbm.26782 [pubmed.ncb…lm.nih.gov], [orca.cardiff.ac.uk]

Jafarian, A., Assem, M. K., Kocagoncu, E., Lanskey, J. H., Fye, H., Williams, R.,…& Rowe, J.B. (2025). Neurophysiological progression in Alzheimer’s disease: insights from dynamic causal modelling of longitudinal magnetoencephalography. Human Brain Mapping, 46(8), e70234.

Jutten, R. J., Peeters, C. F. W., Leijdesdorff, S. M. J., Visser, P. J., Maier, A. B., Terwee, C. B., Scheltens, P., & Sikkes, S. A. M. (2017). Detecting functional decline from normal aging to dementia: Development and validation of a short version of the Amsterdam IADL Questionnaire. *Alzheimer’s & Dementia: Diagnosis*, Assessment & Disease Monitoring, 8, 26–35. 10.1016/j.dadm.2017.03.002

Kim CK, Lee YR, Ong L, Gold M, Kalali A, Sarkar J. Alzheimer’s Disease: Key Insights from Two Decades of Clinical Trial Failures. Journal of Alzheimer’s Disease. 2022;87(1):83–100. 10.3233/JAD-215699

Kocagoncu, E., Nesbitt, D., Emery, T., Hughes, L., Henson, R.H., Cam-CAN & Rowe, J.B. (2022). Neurophysiological and brain structural markers of cognitive frailty differ from Alzheimer’s disease. Journal of Neuroscience, 42(7), 1362–1373. 10.1523/JNEUROSCI.0697-21.2021

Kocagoncu, E., Quinn, A., Firouzian, A., Cooper, E., Greve, A., Gunn, R.,…& Deep and Frequent Phenotyping Study Team. (2020). Tau pathology in early Alzheimer’s disease is linked to selective disruptions in neurophysiological network dynamics. Neurobiology of Aging, 92, 141–152. 10.1016/j.neurobiolaging.2020.03.009

Koychev, I., Lawson, J., Chessell, T., Mackay, C., Gunn, R., Sahakian, B., Rowe, J.B., Thomas, A.J., Rochester, L.R., Chan, D., Tom, B., malhotra, P., Ballard, C., Chessell, I., Ritchie, C.W., Raymont, V., Leroi, I., Lengyel, I., Murray, M., Thomas, D.L., Gallacher, J. & Lovestone, S. (2019). Deep and Frequent Phenotyping study protocol: an observational study in prodromal Alzheimer’s disease. Bmj Open, 9(3), e024498. 10.1136/bmjopen-2018-024498

Lanskey, J. H., Jafarian, A., Karadag, M., Kocagoncu, E., Williams, R. S., Yadav, P., Quinn, A. J., Pitt, J., Thayanandan, T., Lowe, S., Perkinton, M., Timmers, M., Raymont, V., Singh, K. D., Woolrich, M., Nobre, A. C., Henson, R. N., & Rowe, J. B. (2025a). Alzheimer’s disease and its progression reduce pyramidal cell gain and connectivity. Alzheimer’s & Dementia, 21(10), e70805. 10.1002/alz.70805 [pubmed.ncb…lm.nih.gov], [pure-oai.bham.ac.uk]

Lanskey, J. H., Jafarian, A., Hughes, L. E., Karadag, M., Kocagoncu, E., Rouse, M. A., Adams, N. E., Naessens, M., Raymont, V., Woolrich, M., Singh, K. D., Henson, R. N., & Rowe, J. B. (2025b). Alzheimer’s disease and memantine effects on NMDA-receptor blockade: Non-invasive in vivo insights from magnetoencephalography. Molecular Psychiatry. Advance online publication. 10.1038/s41380-025-03288-3

Lanskey, J.H., Kocagoncu, E., Quinn, A.J., Cheng, Y.J., Karadag, M., Pitt, J., Lowe, S., Perkinton, M., Raymont, V., Singh, K.D., Woolrich, M., Nobre, A.C., Henson. R.H., Rowe, J. B & the NTAD study group. (2022). New Therapeutics in Alzheimer’s Disease longitudinal cohort study (NTAD): study protocol. BMJ open, 12(12), e055135. 10.1136/bmjopen-2021-055135

Malpetti, M., Chouliaras, L., Murley, A. G., Swann, P., Coyle-Gilchrist, I., Bevan-Jones, W. R.,…& Rowe, J. B. (2025). Clinicopathological and prognostic correlations of plasma biomarkers in neurodegenerative diseases. medRxiv, 2025-12. 10.64898/2025.12.07.25341788

Malpetti, M., Jones, P. S., Cope, T. E., Holland, N., Naessens, M., Rouse, M. A., Rittman, T., Savulich, G., Whiteside, D. J., Street, D., Fryer, T. D., Hong, Y. T., Milicevic Sephton, S., Aigbirhio, F. I., O’Brien, J. T., & Rowe, J. B. (2023). Synaptic loss in frontotemporal dementia revealed by [11C]UCB-J positron emission tomography. Annals of Neurology, 93(1), 142–154. 10.1002/ana.26543

Mandal, P. K., Banerjee, A., Tripathi, M., & Sharma, A. (2018). A comprehensive review of magnetoencephalography (MEG) studies for brain functionality in healthy aging and Alzheimer’s disease (AD). Frontiers in Computational Neuroscience, 12, Article 60. 10.3389/fncom.2018.00060

Mansur, A., Rabiner, E. A., Comley, R. A., Lewis, Y., Middleton, L. T., Huiban, M., Passchier, J., Tsukada, H., Gunn, R. N., & the MIND-MAPS Consortium. (2020). Characterization of 3 PET tracers for quantification of mitochondrial and synaptic function in healthy human brain: 18F-BCPP-EF, 11C-SA-4503, and 11C-UCB-J. Journal of Nuclear Medicine, 61(1), 96–103. 10.2967/jnumed.119.228080

McNabb, C. B., Driver, I. D., Hyde, V., Hughes, G., Chandler, H. L., Thomas, H., Allen, C., Messaritaki, E., Hodgetts, C. J., Hedge, C., Engel, M.,…Jones, D.K. (2025). WAND: A multi-modal dataset integrating advanced MRI, MEG, and TMS for multi-scale brain analysis. Scientific Data, 12(1), Article 220. 10.1038/s41597-024-04154-7

McKhann, G. M., Knopman, D. S., Chertkow, H., Hyman, B. T., Jack Jr, C. R., Kawas, C. H.,…& Phelps, C.H. (2011). The diagnosis of dementia due to Alzheimer’s disease: recommendations from the National Institute on Aging-Alzheimer’s Association workgroups on diagnostic guidelines for Alzheimer’s disease. Alzheimer’s & Dementia, 7(3), 263–269.

Mecca, A. P., O’Dell, R. S., Sharp, E. S., Godek, T. A., Naganawa, M., Toyonaga, T., Zhao, W., Hicks, K., Perez-Santiago, J., Klunk, W. E., Mathis, C. A., van Dyck, C. H., Carson, R. E., & Chen, M. K. (2022). Synaptic density and cognitive performance in Alzheimer’s disease: A PET imaging study with [11C]UCB-J. Alzheimer’s & Dementia, 18(12), 2527–2536. 10.1002/alz.12582

Mioshi, E., Dawson, K., Mitchell, J., Arnold, R., & Hodges, J. R. (2006). The Addenbrooke’s Cognitive Examination Revised (ACE-R): A brief cognitive test battery for dementia screening. International Journal of Geriatric Psychiatry, 21(11), 1078–1085. 10.1002/gps.1610

Morris, J. C. (1993). The Clinical Dementia Rating (CDR): Current version and scoring rules. Neurology, 43(11), 2412–2414. 10.1212/WNL.43.11.2412-a

Nabulsi NB, Mercier J, Holden D, et al. Synthesis and Preclinical Evaluation of 11C UCB J as a PET Tracer for Imaging the Synaptic Vesicle Glycoprotein 2A in the Brain. Journal of Nuclear Medicine. 2016;57(5):777 to 784. 10.2967/jnumed.115.168179

Nakamura, A., Cuesta, P., Kato, T., Arahata, Y., Iwata, K., Yamagishi, M., Kuratsubo, I., Kato, K., Bundo, M., Diers, K., Fernández, A., Maestú, F., & Ito, K. (2017). Early functional network alterations in asymptomatic elders at risk for Alzheimer’s disease. Scientific Reports, 7(1), Article 6517. 10.1038/s41598-017-06876-8

Nelson HE, & Willison JR. National adult reading test (NART): test manual. 2nd edn, 1991.

Niso, G., Gorgolewski, K. J., Bock, E., Brooks, T. L., Flandin, G., Gramfort, A., Henson, R. N., Jas, M., Litvak, V., Moreau, J. T., Oostenveld, R., Schoffelen, J.-M., Tadel, F., Wexler, J., & Baillet, S. (2018). MEG-BIDS, the brain imaging data structure extended to magnetoencephalography. Scientific Data, 5, 180110. 10.1038/sdata.2018.110

Niso, G., Rogers, C., Moreau, J. T., Chen, L. Y., Madjar, C., Das, S.,…& Baillet, S. (2016). OMEGA: the open MEG archive. Neuroimage, 124, 1182–1187.

O’Dell, R. S., Mecca, A. P., Chen, M. K., Naganawa, M., Toyonaga, T., Godek, T. A., Harris, J. E., Bartlett, H. H., Zhao, W., Nabulsi, N. B., Wyk, B. C., Varma, V. R., McGlinchey, R. E., Pietrzak, R. H., Carson, R. E., van Dyck, C. H., & Hannestad, J. (2021). Association of Aβ deposition and regional synaptic density in early Alzheimer’s disease: A PET imaging study with [11C]UCB-J. Alzheimer’s Research & Therapy, 13(1), 11. 10.1186/s13195-020-00742-y

Perry, A., Hughes, L.E., Adams, N., Naessens, M., Murley, A.G., Rouse, M.A., Street, D., Jones, P.S., Cope, T.E., Kocagoncu, E. & Rowe, J.B. (2022). The neurophysiological effect of NMDA-R antagonism of frontotemporal lobar degeneration is conditional on individual GABA concentration. Translational Psychiatry, 12(1), 348. 10.1038/s41398-022-02114-6

Puttaert, D., Coquelet, N., Wens, V., Peigneux, P., Fery, P., Rovai, A.,…& De Tiège, X. (2020). Alterations in resting-state network dynamics along the Alzheimer’s disease continuum. Scientific reports, 10(1), 21990. 10.1038/s41598-020-76201-3

Quinn, A. J., van Es, M. W. J., Gohil, C., & Woolrich, M. W. (2024). *The osl-ephys Python package for the analysis of electrophysiological data* (Version 2) [Computer software]. Zenodo. 10.5281/zenodo.6875059

Randolph, C., Tierney, M. C., Mohr, E., & Chase, T. N. (1998). The Repeatable Battery for the Assessment of Neuropsychological Status (RBANS): Preliminary clinical validity. Journal of Clinical and Experimental Neuropsychology, 20(3), 310–319. 10.1076/jcen.20.3.310.823

Rosner, B. (1983). Percentage Points for a Generalized ESD Many-Outlier Procedure. Technometrics 25(2), pp. 165–172.

Rowe, C. C., Doré, V., Jones, G., Baxendale, D., Mulligan, R. S., Bullich, S., Stephens, A.W., de Santi, S., Masters, C.L., Dinkelborg, L., & Villemagne, V. L. (2017). 18F-Florbetaben PET beta-amyloid binding expressed in Centiloids. European journal of nuclear medicine and molecular imaging, 44(12), 2053–2059. 10.1007/s00259-017-3749-6

Sabri, O., Sabbagh, M. N., Seibyl, J., Barthel, H., Akatsu, H., Ouchi, Y.,…& Florbetaben Phase 3 Study Group. (2015). Florbetaben PET imaging to detect amyloid beta plaques in Alzheimer’s disease: phase 3 study. Alzheimer’s & dementia, 11(8), 964–974. 10.1016/j.jalz.2015.02.004

Sami, S., Williams, N., Hughes, L. E., Cope, T. E., Rittman, T., Coyle-Gilchrist, I. T. S., Henson, R. N., & Rowe, J. B. (2018). Neurophysiological signatures of Alzheimer’s disease and frontotemporal lobar degeneration: Pathology versus phenotype. Brain, 141(8), 2500–2510. 10.1093/brain/awy180

Selkoe, D. J. (2002). Alzheimer’s disease is a synaptic failure. Science, 298(5594), 789–791. 10.1126/science.1074069

Shafto, M.A., Tyler, L.K., Dixon, M., Taylor, J.R., Rowe, J.B., Cusack, R., Calder, A.J., Marslen-Wilson, W.D., Duncan, J., Dalgleish, T., Henson, R.N., Brayne, C., Cam-CAN & Matthews, F.E. (2014). The Cambridge Centre for Ageing and Neuroscience (Cam-CAN) study protocol: a cross-sectional, lifespan, multidisciplinary examination of healthy cognitive ageing. BMC Neurology, 14(204). 10.1186/s12883-014-0204-1

Sitnikova, T. A., Hughes, J. W., Ahlfors, S. P., Woolrich, M. W., & Salat, D. H. (2018). Short timescale abnormalities in the states of spontaneous synchrony in the functional neural networks in Alzheimer’s disease. NeuroImage: Clinical, 20, 128–152. 10.1016/j.nicl.2018.05.028

Taulu, S., & Simola, J. (2006). Spatiotemporal signal space separation method for rejecting nearby interference in MEG measurements. Physics in Medicine and Biology, 51(7), 1759–1768. 10.1088/0031-9155/51/7/008

Taylor, J.R., Williams, N., Cusack, R., Auer, T., Shafto, M.A., Dixon, M., Tyler, L.K., Cam-CAN, Henson, R.N. (2017). The Cambridge Centre for Ageing and Neuroscience (Cam-CAN) data repository: Structural and functional MRI, MEG, and cognitive data from a cross-sectional adult lifespan sample. NeuroImage, 144, 262–269. 10.1016/j.neuroimage.2015.09.018.

Terry, R. D., Masliah, E., Salmon, D. P., Butters, N., DeTeresa, R., Hill, R., Hansen, L. A., & Katzman, R. (1991). Physical basis of cognitive alterations in Alzheimer’s disease: Synapse loss is the major correlate of cognitive impairment. Annals of Neurology, 30(4), 572–580. 10.1002/ana.410300410

Toyonaga, T., Smith, L. M., Finnema, S. J., Gallezot, J.-D., Naganawa, M., Bini, J., Mulnix, T., Cai, Z., Ropchan, J., Huang, Y., Strittmatter, S. M., & Carson, R. E. (2019). *In vivo* synaptic density imaging with 11C-UCB-J detects treatment effects of saracatinib in a mouse model of Alzheimer disease. Journal of Nuclear Medicine, 60(12), 1780–1786. 10.2967/jnumed.118.223867

Trubshaw, M., Kohl, O., Gohil, C., van Es, M. W., Quinn, A. J., Yoganathan, K., Edmond, E., Proudfoot, M., Zokaei, N., Raymont, V., Pitt, J., Singh, K. D., Woolrich, M. W., Henson, R. N., Hughes, L. E., & Rowe, J. B. (2025). Divergence of cortical neurophysiology across different neurodegenerative disorders compared to healthy ageing. Progress in Neurobiology, 243, 102865.

Spielberger, C. D., Gorsuch, R. L., & Lushene, R. E. (1970). Manual for the State-Trait Anxiety Inventory. Consulting Psychologists Press.

Vaghari, D., Bruna, R., Hughes, L. E., Nesbitt, D., Tibon, R., Rowe, J. B., Maestu, F. & Henson, R. N. (2022). A multi-site, multi-participant magnetoencephalography resting-state dataset to study dementia: The BioFIND dataset. Neuroimage, 258, 119344. 10.1016/j.neuroimage.2022.119344

Villeneuve, S., Poirier, J., Breitner, J.C., Tremblay-Mercier, J., Remz, J., Raoult, J.M., Yakoub, Y., Gallego-Rudolf, J., Qiu, T., Fajardo Valdez, A. & Mohammediyan, B. (2025). The PREVENT-AD cohort: Accelerating Alzheimer’s disease research and treatment in Canada and beyond. Alzheimer’s & Dementia, 21(10), p.e70653.

Wechsler D. Wechsler Adult Intelligence Scale - Revised. San Antonio, TX: Psychological Corporation, 1981.

Wechsler D. Wechsler Memory Scale - Third UK Edition. London: Harcourt Assessment, 1999.

Williams RS, Naessens M, Todd EG, Durcan R, Whiteside DJ, Lanskey J, Jafarian A, Friston K, Hughes LE, Rowe JB (2025). GABAergic regulation of action-outcome priors in conditions associated with frontotemporal degeneration. medRxiv. 10.1101/2025.10.22.25338552

